# Rational design of respiratory syncytial virus dimeric F-subunit vaccines in protein and mRNA forms

**DOI:** 10.1101/2025.05.31.655981

**Authors:** Jing Li, Xuehui Ma, Zepeng Xu, Wenjing Guo, Ruchao Peng, Yuqin Zhang, Yumin Meng, Jianrui Zhao, Qiyue Wang, Shuang Li, Jiaqin Chen, Yuxin Guo, Xuancheng Lu, Qingling Wang, Yushuang Guo, Meng-ao Jia, Yan Li, Yanfang Zhang, Shihua Li, Pei Du, Qihui Wang, George Fu Gao, Jianxun Qi

**Affiliations:** Laboratory of Pathogen Microbiology and Immunology, Institute of Microbiology, Chinese Academy of Sciences, Beijing 100101, China; Beijing Life Science Academy, Beijing 102209, China; Zhejiang University School of Medicine, Hangzhou, Zhejiang 310058, China; Faculty of Health Sciences, University of Macau, Macau SAR 999078, China; University of Chinese Academy of Sciences, Beijing 101408, China; Department of Biochemistry and Biophysics, Perelman School of Medicine, University of Pennsylvania, Philadelphia 19104, PA, USA; School of Laboratory Medicine and Life Sciences, Wenzhou Medical University, Wenzhou, Zhejiang 325035, China; National Key Laboratory of Intelligent Tracking and Forecasting for Infectious Diseases (NITFID), Chinese Center for Disease Control and Prevention, Beijing, 102206, China; Shanxi Natural Carbohydrate Resource Engineering Research Center, College of Food Science and Technology, Northwest University, Xi’an 710069, China; Key Laboratory of Molecular Genetics, China National Tobacco Corporation, Guizhou Institute of Tobacco Science, Guiyang, Guizhou, 550083, China

## Abstract

**Background:** Respiratory syncytial virus (RSV) poses a significant public health threat, particularly to children and the elderly. Two protein-based vaccines and one mRNA vaccine have been approved, all targeting the prefusion conformation of the fusion (F) glycoprotein. However, it has been reported that the F protein transitions to the post-fusion state during storage, resulting in a reduction of the vaccines’ immunogenicity.

**Methods:** In this study, we engineered novel pre-F-based antigens to preserve pre-F-specific immunodominant epitopes while eliminating sub-potent ones. Based on this, we constructed a series of single-chain dimers and selected the one with the highest expression yield and melting temperature (*T*_m_). Next, we created a heterodimer, scDimer AB. Structural and protein characterization analyses were conducted to verify our design. All monomeric and scDimer antigens were used to immunize rodent models. Additionally, we prepared the antigens in mRNA form and immunized BALB/c mice. Finally, we combined both antigen forms, administering intramuscular mRNA priming followed by intranasal protein delivery in mice. In all immunization strategies, viral challenges were performed in animals to evaluate the immunologic protective effects.

**Findings:** Through rational design, we developed a monomeric and two single-chain dimeric (scDimer) proteins with the expected characteristics, including complete II, V, and Ø epitopes and a partial III epitope. The scDimers elicited stronger binding and neutralizing antibody responses in rodent models compared to the monomer, and they also boosted T cell responses when combined with appropriate adjuvants. After three doses of scDimer immunization, live RSV was barely detectable in the tissues of infected animals. The copies of RNA encoding N-gene were significantly reduced in the immunized groups compared to the PBS-injected control groups. We also engineered mRNA versions of the antigens and verified their protective efficacy in mice. Notably, there were no significant differences between intranasal boost and intramuscular two doses after RSV challenged, suggesting that intranasal boost provided equivalent protection to intramuscular vaccination and could reduce the risk of vaccine-enhanced disease (VED) potentially.

**Interpretation:** The scDimer-based RSV vaccines effectively protected rodents from RSV infections, highlighting their clinical potential. Our antigen design removed certain suboptimal epitope regions, enhancing the efficiency of antigen presentation and increasing the proportion of the most potent pre-F-specific neutralizing antibodies. This approach provides a novel perspective for future vaccine design.

**Funding:** National Key R&D Program of China, National Science Foundation of China, Young Scientists in Basic Research, Chinese Academy of Sciences, and Special Program of China National Tobacco Corporation.

**RESEARCH IN CONTEXT:** *Evidence before this study:* The licensed respiratory syncytial virus (RSV) vaccines were based on RSV pre-F structure. We conducted a PubMed search for articles published in English from database’s inception to March 31, 2025, using the search terms “RSV” (interchanged with “respiratory syncytial virus”), “pre-F” (interchanged with “prefusion F”) and “vaccine”. Our search revealed that most of the articles focus on combining pre-F with RSV G protein or other proteins to form viral-like particles (VLPs) or nanoparticles. The second-largest category of articles examined the antibodies level elicited by the pre-F antigens in animal model or some special cohorts. Other studies addressed topics such as optimization of injection dosage and immunization strategies, methodologies for detecting pre-F-specific antibodies, and the development of mRNA-LNP vaccines. Notably, only 4 articles described novel designs of RSV pre-F antigens, all of which were trimeric and based exclusively on the RSV A subtype F protein sequence.

*Added value of this study:* To our knowledge, our design is the first single-chain dimer version of RSV pre-F antigen. The engineered antigens demonstrated high yield, thermostability, and immunogenicity comparable to that of DS-Cav1. Additionally, we developed the first chimeric RSV-antigen covering both subtype A and B, thereby broadening the potential protective spectrum. We further validated the immunogenicity of our designs in mRNA form and as a protein-mRNA combination. This research emphasizes the potential of constructing and combining minimized, stable antigen modules, offering a novel approach for future vaccine development.

*Implications of all the available evidence:* In this study, we present a novel approach for constructing and combining minimized, stable antigen modules for future vaccine design. The strong immunogenicity of our antigens underscores their potential for clinical application. Furthermore, given the structural similarities between the RSV antigen and those of other *Pneumoviruses*, such as human metapneumovirus (hMPV), our design strategy could be extended to the development of multivalent vaccines.

## INTRODUCTION

Respiratory syncytial virus (RSV) infection can lead to severe morbidity and mortality, particularly among children under 5 years old^1^ and the elderly.^2^ In 2019, over 33 million children were infected with RSV, resulting in over 10% hospitalization and more than 26,000 deaths. Infants under six months face an even greater risk, with 1⋅4 million hospital admissions and over 13,000 deaths.^3^ Among adults over 65 years old in industrialized countries, 1⋅5 million RSV infection cases were reported in 2015, with a fatality rate of approximately 5%.^4^ Low- and middle-income countries (LMICs) have a higher burden of RSV infections. As the global population ages, the societal impact of RSV is expected to increase. While two monoclonal antibodies targeting the fusion (F) glycoprotein have been approved for prophylaxis in newborns,^5,6^ vaccines remain the most effective preventive measure.

RSV is a single-stranded negative-sense RNA virus belonging to the *Orthopneumovirus* genus, *Pneumoviridae* family, and is classified into subtypes A and B based on genome sequences.^7^ Epidemiological investigations have revealed that subtype A is more prevalent globally, accounting for about 75% of cases in most years.^8^ The RSV genome, which is 15⋅2 kb in length, encodes 11 viral proteins. The conserved F glycoprotein plays a critical role in mediating fusion between the virus and host cell membranes, facilitating viral entry. During the virus-cell fusion process, the F protein undergoes a conformational change from its pre-fusion (pre-F) to post-fusion (post-F) state. The pre-F induces stronger neutralizing antibody responses compared to the post-F,^9^ and the pre-F-specific Ø and V epitopes are responsible for eliciting most potent neutralizing antibodies.^10^ Thus, stabilizing the pre-F conformation is a primary goal of most RSV vaccine development strategies. DS-Cav1, a representative pre-F stabilized by introducing a disulfide bond and two cavity-filling substitutions, is adopted in the approved GSK protein-based vaccine, Arexvy.^10,11^ An enhanced version, DS2, with additional disulfide bonds and substitutions, was also adopted in Moderna’s licensed mRNA vaccine (mRNA-1345, mRESVIA).^12^ In addition, Pfizer has developed pre-F stabilized proteins using different substitutions (Abrysvo), which have also been approved by the FDA.^13^ However, it has been reported that DS-Cav1 may transit to post-F upon storage.^14,15^

While the F protein is generally conserved across RSV subtypes A and B (with 89% sequence identity), the epitope Ø shows divergence, inducing subtype-specific antibodies.^16^ To achieve broad protection, Pfizer mixes F proteins from both subtypes to produce a bivalent vaccine.^13^ However, mixed vaccines might require higher dosages and could potentially induce non-neutralizing antibodies. In summary, optimizing vaccine stability, efficiency, and cost-effectiveness is crucial for rational design of RSV vaccines.

## Methods

### Study design

In this antigenic study, we designed monomer A, an antigen based on RSV subtype A F protein, and determined its crystal structure. To enhance immunogenicity, we created a single polypeptide construct where two monomers are tandemly linked but folds independently (single-chain dimer, scDimer). To expand the protection spectrum, we then designed monomer B based on subtype B F protein and fused it with monomer A to form a single-chain heterodimer AB. After proteins characterization, we immunized rodents with three doses of the recombinant antigen at three-week intervals.

To assess the immunoprotection of the two scDimer antigens, a nasal virus challenge was performed three weeks after the 3^rd^ immunization. Additionally, the two scDimer antigens were administered to mice in mRNA form, with two doses given at two-week intervals. Finally, a combination of protein and mRNA antigen formulations, as well as other immunization protocols, were evaluated to expand the scope of our design’s application. All animal experiments were approved by the Ethics Committee IMCAS and conducted in accordance with the guidelines outlined in the Guide for the Care and Use of Laboratory Animals.

### Procedures

#### Protein expression, purification, and crystallization

For each construct, the signal peptide sequence of the RSV F protein (residues 1-25) was added to the protein’s N terminus for secretion, and a hexa-His tag was attached at the C terminus for efficient purification. The purified monomer A and scDimer AA were deglycosylated by PNGase F at room temperature for 3 hours. Crystals of the deglycosylated proteins were grown by sitting-drop method at 18 °C. X-ray diffraction data were collected at Shanghai Synchrotron Radiation Facility (SSRF) BL02U1 and BL19U1. All structural visualization were generated using Pymol software (https://pymol.org/).

#### Protein characteristics assay

Surface plasmon resonance (SPR) was employed to analyze the binding kinetics between the protein and monoclonal antibodies (MAbs). The Prometheus NT.48 instrument and nano-DSC were used to evaluate the melting temperature (*T*m) of the proteins. The stability of key epitopes was evaluated using Octet Red96. Expression yields were determined by Jess^TM^ ProteinSimple. Detailed methods can be found in the Supplementary materials.

#### Animal immunization and viral challenge

Specific pathogen-free (SPF) female BALB/c mice or cotton rats, aged 6 to 8 weeks, were used for immunization. They were housed under SPF conditions in the laboratory animal facilities of the Institute of Microbiology, Chinese Academy of Sciences (IMCAS), and the Chinese Center for Disease Control and Prevention (China CDC), Laboratory Animal Center. RSV challenge experiments were performed in an animal biosafety level 2 (ABSL2) facility at China CDC, Laboratory Animal Center. For protein-antigen experiments, rodents were intramuscularly injected three times at a 3-week interval. Peripheral blood was collected two days before the 2^nd^ and the 3^rd^ immunization or two weeks after the 3^rd^ immunization, for serum separation. In mRNA- and mRNA-protein combined antigen studies, mice were immunized twice at a 2-week interval. Peripheral blood was collected two days before the 2^nd^ immunization or two weeks after the 2^nd^ immunization, and serum was separated. The nasal virus challenge was performed on the 7^th^ day following the last time of blood collection. Five days later, the animals were executed, and the tissues were collected.

#### Titer assay, cytokine assay, and tissue stain

The serum samples were analyzed to determine the titers of binding and neutralizing antibodies. Lung, trachea, and nasal tissues were used to quantify the loading virus titer by qRT-PCR and culture. Parts of lung tissue were assessed for the cytokine level by flow cytometry. Other parts of lung tissue samples were stained with H&E. The spleen was used to detect T cell responses by ELISpot.

### Statistical analysis

For qRT-PCR assay, standard curves were generated using known 7 titers of virus N gene amplification and performed in GraphPad Prism (version 9⋅0) by a linear regression. Each titer point was repeated three times. The R^2^ values of the standard curves were greater than 0⋅97.

For the antibody titer assay, each sample was performed with two technical replicates. Data were presented as geometric mean ± 95 % confidence interval (CI). For ICS assays, data are presented as mean ± standard error. ANOVA test was used to assess the difference between groups and *P* values less than 0⋅05 were considered statistically significant. All graphs and statistical analyses were generated using GraphPad Prism version 9⋅0 software.

*K*_D_ values for SPR assays were calculated by the software BIAevaluation Version 4⋅1 (GE Healthcare), using a 1:1 binding model. The values are shown as mean ± SD of two independent experiments. Details can be found in figure legends.

## RESULTS

### Rational design for preserving pre-F epitopes of Ø and V and maintaining their antigenicity

Previous data indicate that epitopes Ø and V on the F protein are the most potent in eliciting neutralizing antibodies against RSV infection, while other epitopes exhibit relatively weaker activity. However, in the post-fusion F structure, epitopes Ø and V undergo substantial conformational changes, sabotaging their neutralizing sensitivity. Thus, we aim to design a vaccine candidate that satisfies two key requirements: preservation of epitopes Ø and V, and stabilization of the prefusion conformation. It has been reported that two refolding regions experience significant conformational change during fusion process (Figure S1A).^10^ Moreover, the C-terminal region of RSV F protein is distant from epitopes Ø and V. We hypothesized that truncation of the C-terminus could inhibit the conformational changes. Based on the structure of DS2 (PDB: 5k6i),^17^ we engineered the RSV F protein that preserves epitopes Ø, V, and II by truncating the C-terminus and preserving the head region containing these epitopes. To prevent undesired inter-chain disulfide bonds, we introduced C149A substitution. During attempts to determine the crystal structure of the new antigen, we encountered difficulties with crystallization. To overcome this, we introduced an additional substitution, G46S inversion, and successfully resolved the structure of monomer A at 2⋅1 Å (Figure 1A, Table 1). Compared with the full-length pre-F (PDB: 5k6i), the prefusion conformation is well maintained, with the root mean square deviation (RMSD) of 0⋅517 Å for 202 C_α_ atoms (Figure S1A). Notably, during antigen production, dimeric antigens were formed by disulfide bonds. To prevent dimer formation, we introduced another mutation C37A. Consistent with our design, monomer A exhibited robust affinities towards monoclonal antibodies (MAbs) targeting epitopes II,^18^ V^19^, and Ø,^20^ with dissociation constant (*K*_D_) values of nanomolar or even picomolar. In contrast, the binding affinity towards an MAb directed at epitope III^21^ was significantly diminished (*K*_D_ > 10 μM), likely due to the partial deletion of this epitope. No detectable binding was observed for MAbs targeting epitopes I^15^ or IV^22^ (Figure 1B, S1B, Table 2).

**Figure 1.**
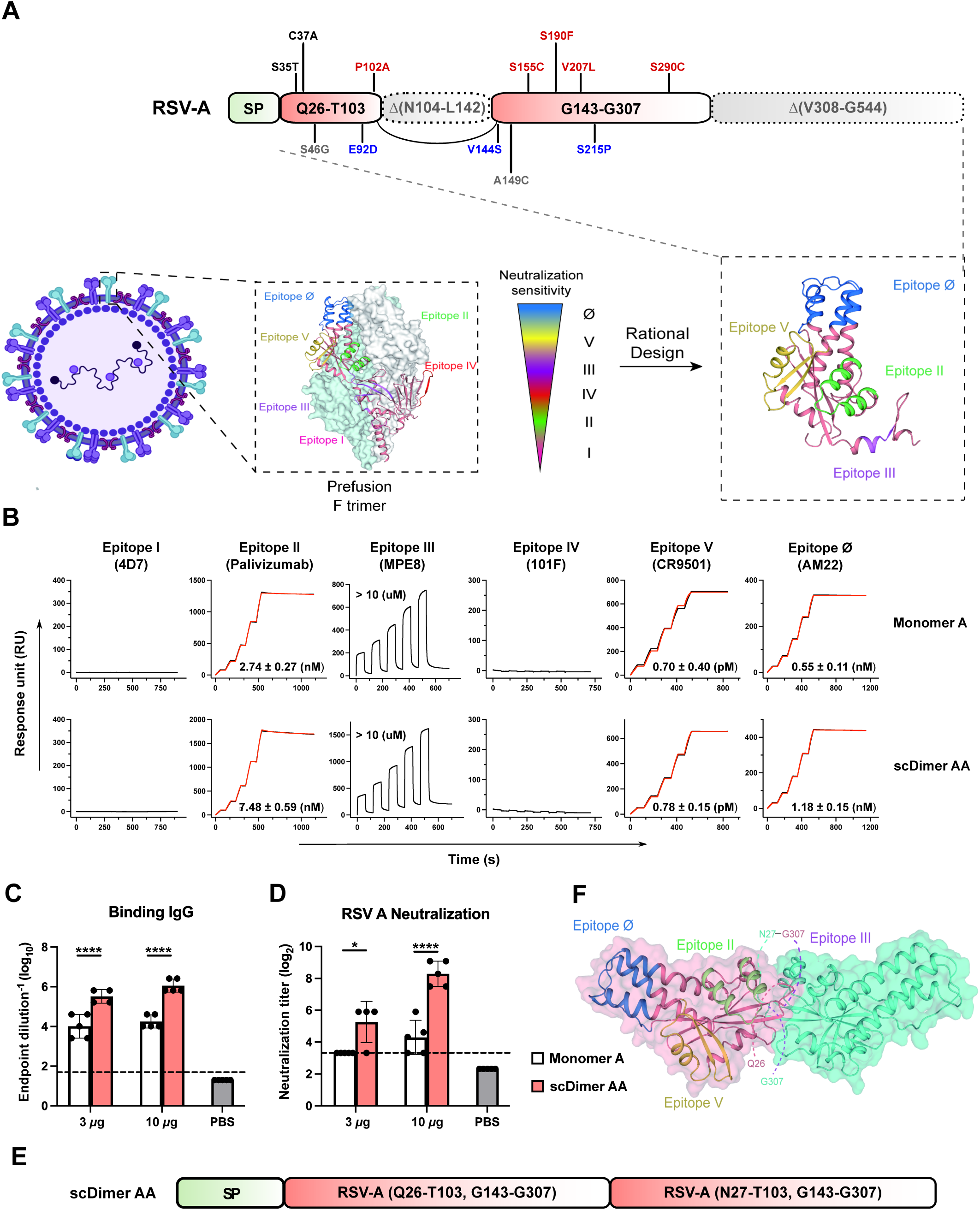
Rational design of RSV F protein monomer A and scDimer AA at pre-fusion state and their immunogenicity assessment. (A) Schematic diagram and design rationale of monomer A. The design of monomer A was outlined (top). Substitutions unique to monomer A and DS2 are labeled in black and gray, respectively. Substitutions shared among DS-Cav1, DS2, and monomer A are colored in red, and those shared between DS2 and monomer A are colored in blue. The dashed squares indicated the deleted residues. In the pre-F structure (PDB: 5k6i), Epitope I, II, III, IV, V, and Ø were labeled in magenta, green, purple, red, yellow, and blue, respectively and their neutralizing sensitivities are indicated. (B) Binding affinities of monomer A (top) and scDimer AA (bottom) with MAbs targeting six epitopes. Actual and fitted curves were represented by red and black lines, respectively. *K*_D_ values were listed as mean ± SD of three experiments. (C) Endpoint titers of immunized antigen-binding IgG in murine sera were measured by ELISA. Groups of mice (n=4∼5) were intramuscularly immunized with 3 or 10 µg of monomer A or scDimer AA at 0, 3, and 6 weeks with Alhydrogel^®^ adjuvant. Serum samples were collected on day 56 (two weeks after the 3^rd^ immunization) and subjected for ELISA test. (D) Neutralization of RSV A2 strain in murine sera on day 56 was assessed. (E) Design of scDimer AA. Two truncated monomers A were tandemly linked, with the latter one starting at N27 residue. (F) Crystal structure of scDimer AA, with epitopes labeled as in panel A. Dotted lines represented areas with uncharacterized structures due to the flexibility of the regions. In (C) and (D), dotted lines indicated the lower limit of detection (LLD). Any value less than the LLD was assigned a value of 1⁄2 the LLD. Data were derived from two technical replicates of a single experiment, with bars indicating mean ± SD. *P* values were determined using Two-way ANOVA (ns indicated *P* > 0⋅05; * indicated *P* < 0⋅05; ** indicated *P* < 0⋅01, *** indicated *P* < 0⋅001, **** indicated *P* < 0⋅0001).

**Table 1.**
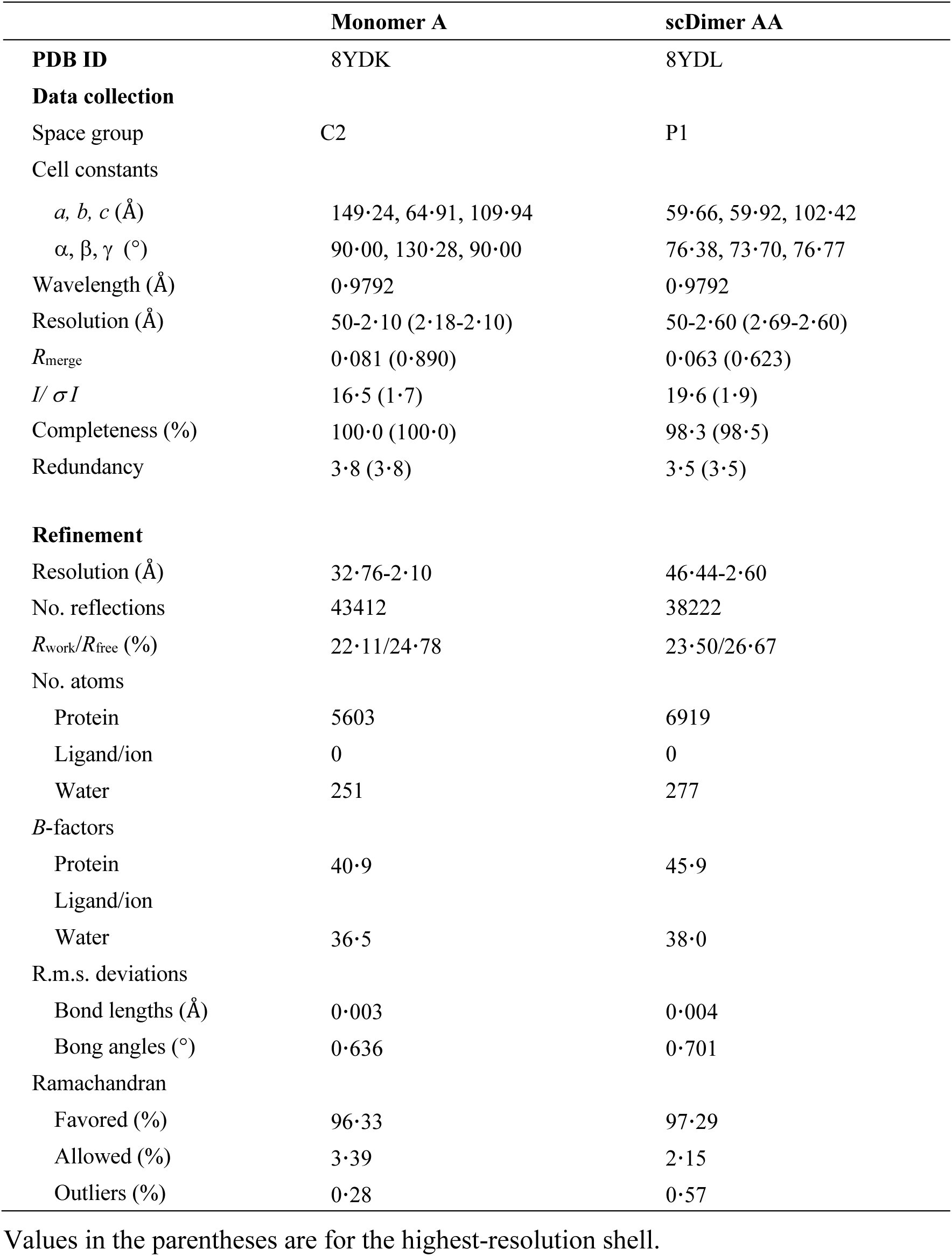
Crystallographic data collection and refinement statistics.

**Table 2.**
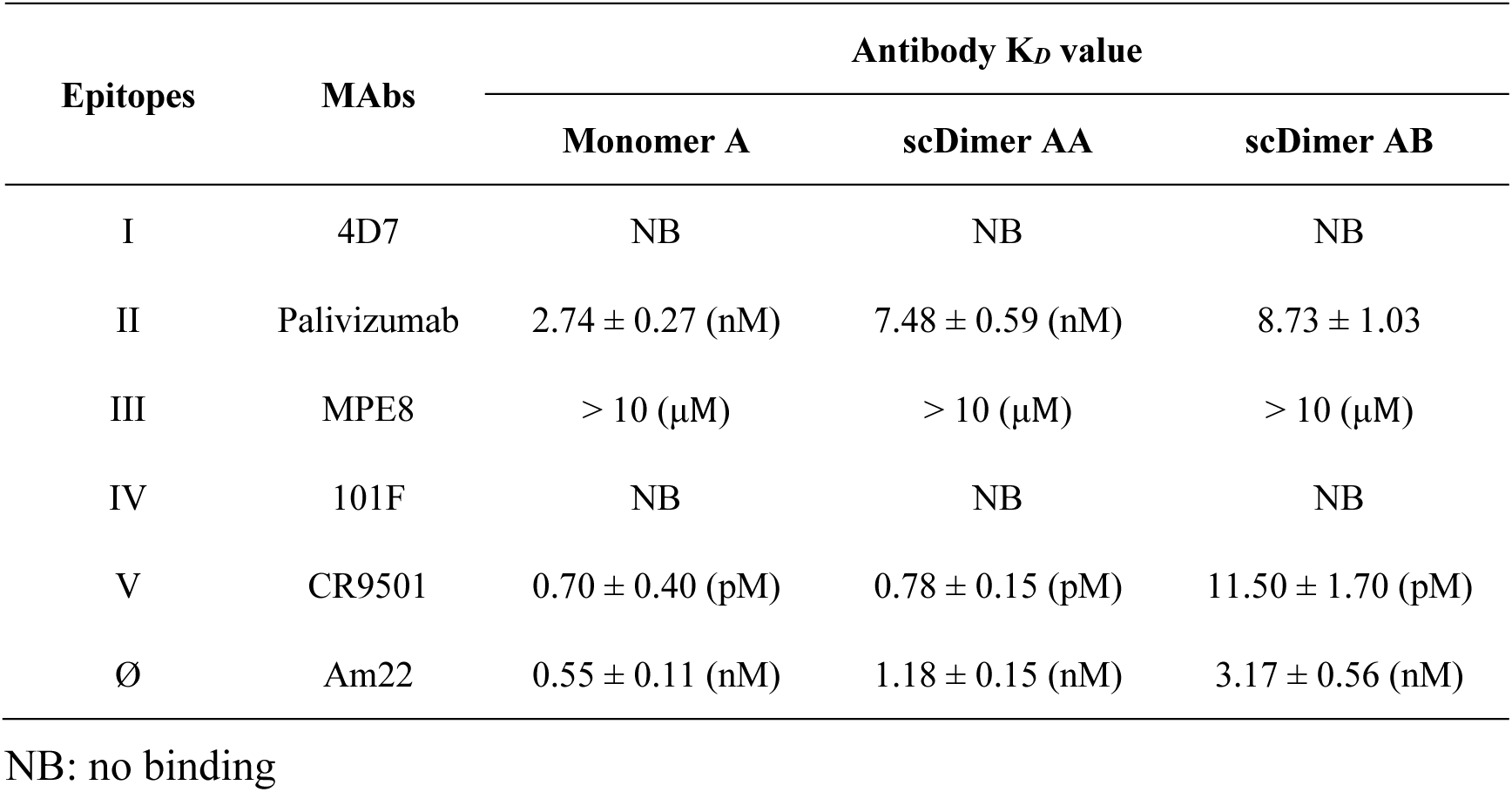
Binding affinities of antigens bound to representative MAbs targeting six major epitopes.

The immunogenicity of the monomer A protein was subsequently evaluated in mice (Figure 1C). Although the immunized sera exhibited apparent antibody responses against the monomer A protein compared to that of the PBS-adjuvant group, there was only a less than 10-fold increase in neutralization efficiency against the RSV A2 strain even after three doses of vaccination (Figure 1C, D, and Figure S4B, C). These results suggest that the monomer A protein may have limited effectiveness for further applications.

### Enhanced immunogenicity and protective efficacy of tandemly repeated antigen dimer

To enhance the immunogenicity of the antigen, we adopted a strategy of tandemly linking two copies of the antigen, following our previous success in vaccines against the severe acute respiratory syndrome coronavirus 2 (SARS-CoV-2) and the monkeypox virus (MPXV).^23,24^ To identify the most stable fused protein, we designed four constructs (Figure S2A), incorporating either an intact or truncated version of epitope III (residue 27-41) between two monomer units (Figure S2A). The yield and thermal stability (*T*m) of these four single-chain dimer (scDimer) constructs were evaluated. Among these constructs, the first construct (named scDimer AA) exhibited the highest yield (> 60 mg/L) and *T*m value (> 79 ℃), which is also the highest yield among the reported constructs,^17,25^, and was selected for further characterization (Figure 1E, S2B, S3A, and B). Notably, the *T*m of scDimer AA was significantly higher than that of DS-Cav1, which has a *T*m of only 51⋅94 ℃ (Figure S3C).

We further evaluated the properties of the scDimer AA. The crystal structure of scDimer AA was resolved at 2⋅6 Å (Table 1), confirming full exposure of epitopes Ø, V, and II on the surface as modeled (Figure 1F). Consistently, this scDimer exhibited comparable binding affinities to the corresponding MAbs as monomer A (Figure 1B). Furthermore, scDimer AA demonstrated remarkable stability. Comparing with the 0 day data, the response retained over 50 % reactivity to antibodies even after 100 days of storage at 4 ℃ (Figure S3D). The Ø and V epitope-reactivity of this protein showed minimal reduction after 4 weeks at 37℃ (Figure S3E), while DS-Cav1 exhibited a significant loss of reactivity under the same storage condition (Figure S3F).

To assess the immunogenicity of scDimer AA, we performed a comparative analysis using BALB/c mice (Figure S4A). Mice vaccinated with scDimer AA elicited significantly stronger antigen-specific antibody responses compared to those vaccinated with monomer A, with antibody titers nearly 100-fold higher across all immunization cycles (Figure 1C and Figure S4B). Additionally, sera from the scDimer AA-immunized group demonstrated an 8- to 500-fold increase in neutralization efficiency against RSV A2 strain compared to the unvaccinated group, and the neutralization levels were also significantly higher than those observed in the monomer A group (Figure 1D and Figure S4C).

We further evaluated the cellular immune efficacy of these antigens adjuvanted with AddaVax^TM^, an adjuvant that is known to elicit stronger cellular immunity compared to Alhydrogel^®^, using the ELISpot assay. Following induction with the monomerA peptide pool, the number of spots was quantitied using CTL. The results demonstrated that AddaVax^TM^-adjuvanted antigens elicited a robust T cell response, with scDimer AA slightly higher than monomer A (Figure S4D). Collectively, these findings suggested that the scDimer design exhibited potency in eliciting both neutralizing antibodies and cellular immunity *in vivo*, and is worth further investigation.

Using the cotton rat as an RSV infection model,^14^ we investigated the protective efficacy of scDimer AA *in vivo* (Figure S5A). Mice injected with a PBS-adjuvant mixture were used as negative control. Compared to the negative control group, all scDimer AA-immunized groups exhibited apparent antibody responses following each immunization (Figure S5B). Notably, after the 3^rd^ immunization, sera from the scDimer AA immunized group exhibited an approximately 100-fold increase in response specific to scDimer AA, which contains the Ø, V, II, and partial III epitopes, compared to the response against post-F which contains epitopes I, II, III and IV. This indicates that the novel antigen primarily elicited antibodies specific to the pre-F Ø and V *in vivo* (Figure S5C). Higher dosage of Alhydrogel^®^-adjuvanted scDimer AA elicits a stronger humoral response against RSV A2 (Figure 2A). In AddaVax^TM^ -adjuvanted groups, while there was no significant difference between 3 µg and 10 µg groups, the 3 µg group on average elicited higher neutralizing antibodies against RSV (Figure 2A, B). Both Alhydrogel^®^- and AddaVax^TM^-adjuvanted groups elicited high titers of neutralizing antibodies against RSV A2 (256-1024) and B9320 strains (256-2048) (Figure 2A, B and Figure S5D, E). After challenging with 1 × 10^5^ TCID_50_ RSV A2 strain, all scDimer AA-immunized groups exhibited reduced viral titers in the lung and trachea, and slightly lower in the nasal tissue (Figure 2C). As the qRT-PCR primer and probe could not distinguish the nucleotide shreds from live viruses, it is suggested that the RNA detected in nasal tissues represents the remains of the challenging virus. In summary, scDimer AA induced a robust immune response that inhibited virus replication, leading to reduced virus titers, particularly in the lower respiratory tract.

**Figure 2.**
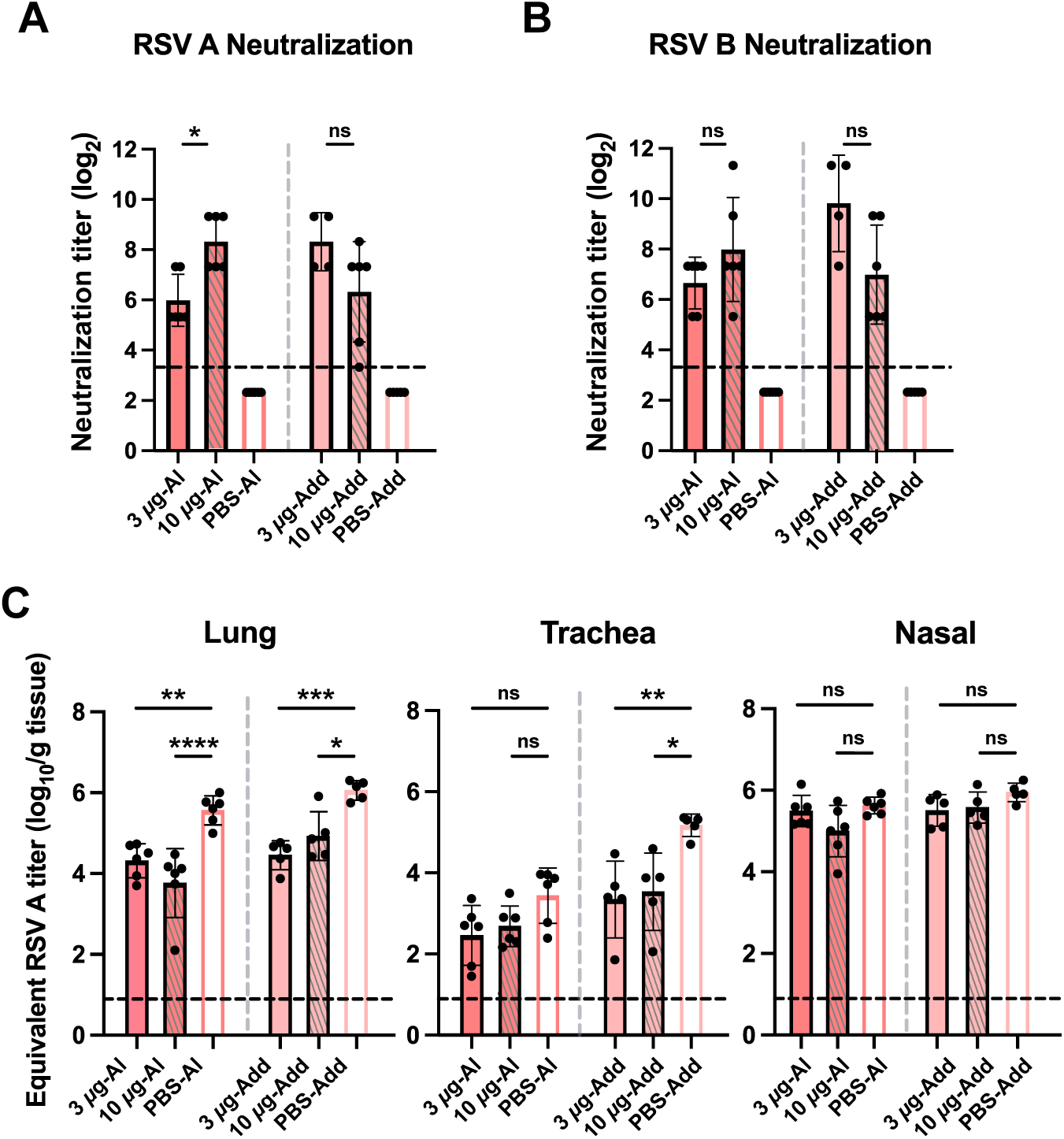
Immunogenicity and protective efficacies of scDimer AA in cotton rats. (A)-(B) Neutralizing titers of cotton rat sera (n=5∼6) against RSV A2 (A) and RSV B9320 (B) strains after the third immunization. (C) Equivalent RSV A titers of lung, trachea, and nasal tissues on 5 dpi. For panels (A) and (B), data were from two technical replicates of a single experiment. For panel (C), data were from three technical replicates of a single experiment. The dotted lines indicate the LLD. Any value less than the LLD was assigned a value of 1⁄2 the LLD. Bars represent mean ± SD. *P* values were analyzed with One-way ANOVA (ns indicated *P* > 0⋅05; * indicated *P* < 0⋅05; ** indicated *P* < 0⋅01, *** indicated *P* < 0⋅001, **** indicated *P* < 0⋅0001).

### Immunogenicity and protective efficacy of the bivalent scDimer design

To broaden the protection spectrum of the antigen, we engineered a chimeric scDimer (scDimer AB) by incorporating antigens from both RSV A and B (RSV B18537) subtypes (Figure 3A, Figure S6A). The scDimer AB exhibited binding characteristics similar to scDimer AA, exhibiting non-dissociating binding patterns with Palivizumab,^18^ CR9501,^19^ and AM22,^20^ a rapid association-dissociation model to MPE8,^21^ and no response to 4D7^15^ or 101F^22^ (Figure S6B, Table 2). The thermostability of scDimer AB was comparable to that of scDmier AA with a *T*m value of 83⋅27 ℃ (Figure S6D). The Ø and V epitope-reactivity retained over 50 % even after 4 weeks of storage at 37 ℃ (Figure S6C). In a cotton rat immunization experiment, a positive group was injected with the same dosage of a DS-Cav1-adjuvant mixture, and a negative control group received a PBS-adjuvant mixture. Similar to scDimer AA, scDimer AB elicited robust antibody responses following each immunization, showing responses comparable to DS-Cav1 (Figure S6E and F). However, unlike the DS-Cav1 injected groups, antibodies induced by scDimer AB predominantly targeted the pre-F conformation (Figure S6F). Like scDimer AA, scDimer AB sera also exhibited high titers of neutralizing antibodies against both RSV A2 (32-4096) and B9320 (64-8192) (Figure 3B, C and Figure S6G, H), comparable to those seen in the DS-Cav1 injected groups.

**Figure 3.**
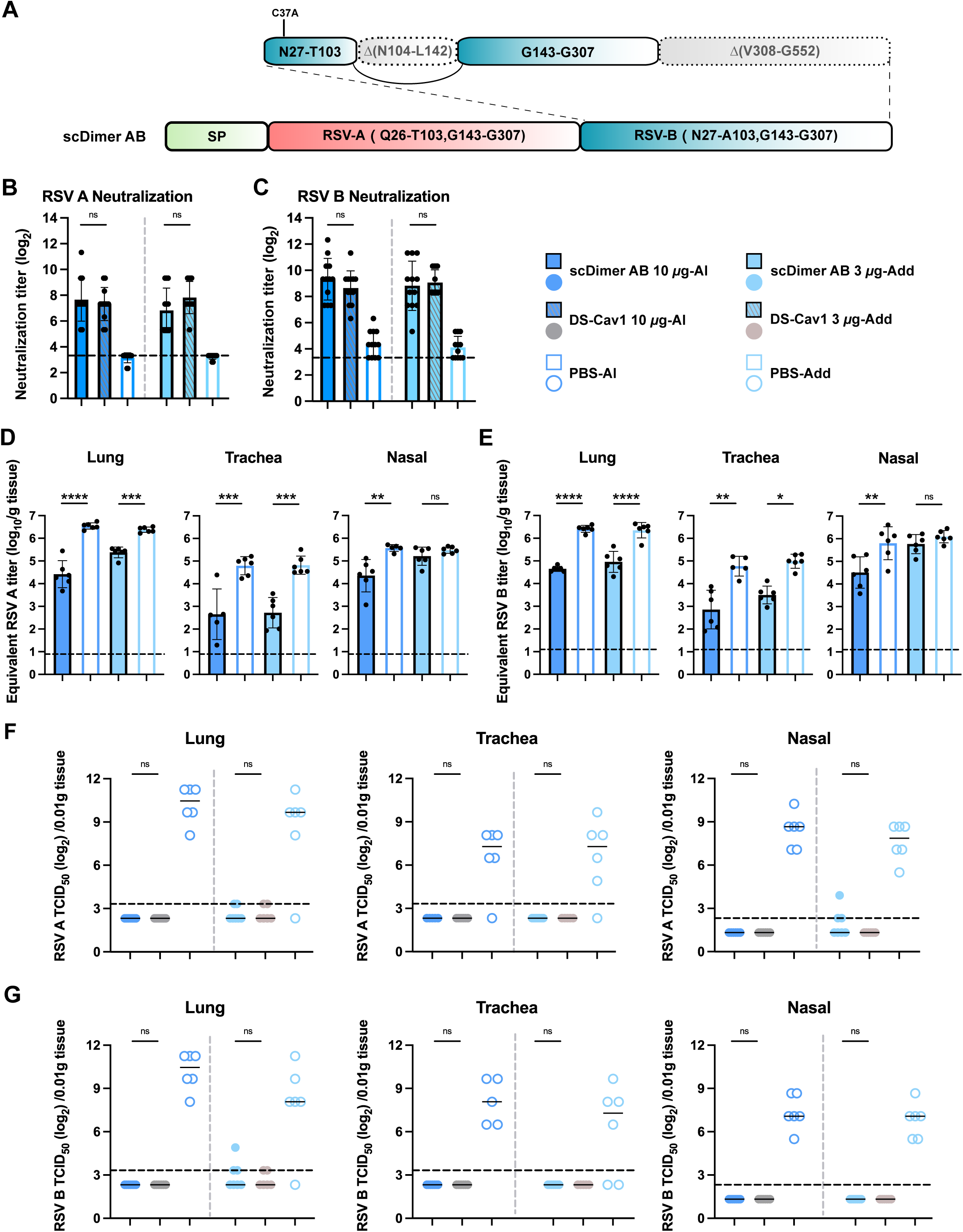
Immunogenicity and protective efficacies of scDimer AB and DS-Cav1 in cotton rats. (A) The schematic diagram of scDimer AB. The RSV-A part is the same as presented in Figure 1A. The deleted portion of RSV B part, compared to the RSV B18537 strain, is indicated by dashed squares. (B)-(C) Neutralizing titers of cotton rat sera (n=12∼14) against RSV A2 (B) and RSV B9320 (C) strains after the third immunization. (D)-(E) Equivalent RSV A2 (n=6∼7) (D) and RSV B9320 (n=6∼7) (E) titers of the lung, trachea, and nasal tissues on 5 dpi. (F)-(G) Cytopathogenic effect (CPE) assays for A2 (n=6∼7) (F) and B9320 (n=6∼7) (G) were performed in HEp-2 cells. For panels (B), (C), (F), and (G), data were from two technical replicates of a single experiment. For panels (D) and (E), data were from three technical replicates of a single experiment. The dotted lines indicate the LLD. Any value less than the LLD was assigned a value of 1⁄2 the LLD. Bars represent mean ± SD. *P* values were analyzed with One-way ANOVA (ns indicated *P* > 0⋅05; * indicated *P* < 0⋅05; ** indicated *P* < 0⋅01, *** indicated *P* < 0⋅001, **** indicated *P* < 0⋅0001).

When challenged with 1 × 10^5^ TCID_50_ RSV A2 or B9320 strain, all vaccinated groups displayed lower viral loads in the lung and trachea compared to the negative control, with a decrease ranging from 31- to 251-folds. In the nasal tissue, the Alhydrogel^®^-adjuvanted group exhibited significantly decreased viral loads (13-fold) (Figure 3D, E). To further validate the protective potential of this vaccine candidate *in vivo*, homogenized tissue samples were co-cultured with single-layer HEp-2 cells in 96-well plates for 4 days to determine virus titer. According to cytopathogenic assay, live RSV was hardly detected in tissues of the immunized groups, whereas in the PBS groups, the average viral titer was 32- to 256-fold higher than those in the immunized groups (Figure 3F, G). Importantly, there was no significant difference between scDimer AB and DS-Cav1 injected groups. These data demonstrated that scDimer AB vaccination could effectively inhibit viral infection and provide protection against both RSV subtypes *in vivo*.

### mRNA-based scDimers also provided effective protection for mice against both RSV subtypes

mRNA vaccines offer significant advantages in stimulating not only humoral but also cellular immunity. Therefore, we also investigated the efficacy of the scDimer constructs in the form of mRNA vaccines. The lipid nanoparticle (LNP)-encapsulated mRNAs encoding scDimer AA and AB were individually prepared (Figure S7A), we also measured the expressions of scDimer AA and AB-mRNA in HEK293 cells. The results demonstrated that both scDimers were effectively expressed between 12-48 hours (Figure S7B). Subsequently, we immunized mice with LNPs (Figure 4A). The two immunized sera revealed comparable high titers of antibodies specific to the respective immunogens (the group of LNP was coated with AB antigen) (Figure 4B), and exhibited robust and comparable neutralizing activities against both RSV strains A2 and B9320 (Figure 4C, D). As anticipated, upon re-stimulation with peptide pools of both subtype A and B, enzyme-linked immune absorbent spot (ELISpot) assays showed a significant increase in the number of IFNγ spots in both groups. Notably, the scDimer AB group exhibited balanced subtype A- and B-specific T cell response, with spots in both groups reaching approximately 400 spots per 0⋅25 million cells (Figure 4E). While the scDimer AB group displayed a higher response towards subtype B than A.

**Figure 4.**
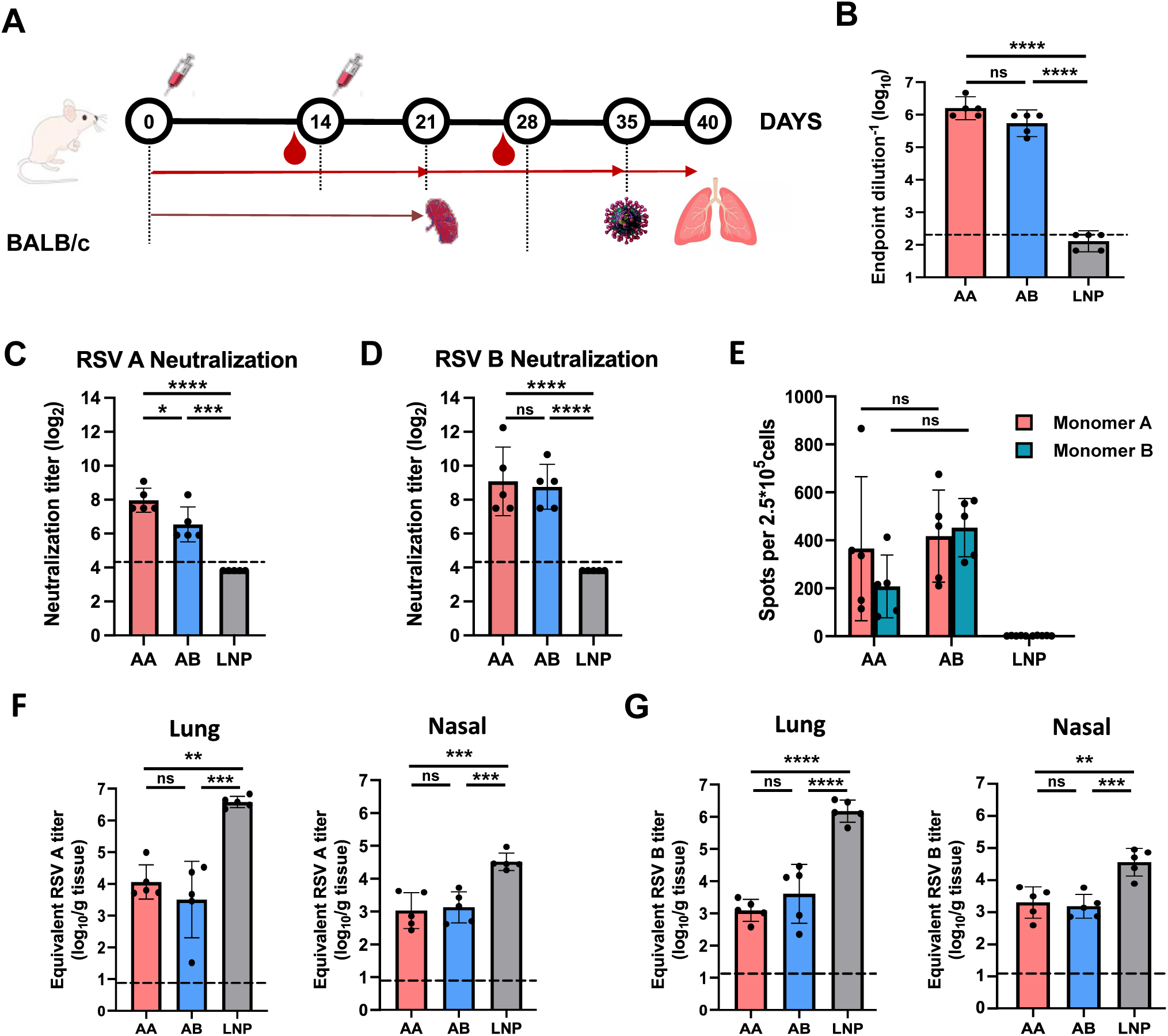
Evaluations of immunogenicity and protective efficacies of the mRNA vaccines encoding scDimers in mice. (A) Procedure for mice immunization and sample collections. Groups of 6 to 8-week-old female BALB/c mice (n=5) were intramuscularly immunized with 5 µg of mRNA-LNP on day 0 and 14. Serum samples were collected on day 12 and 26. The immunized rats were then challenged with intranasal administration of 1×10^6^ TCID_50_ of RSV subtype A2 on day 35. Lung and nasal tissues were harvested 5 days post infection (dpi). Spleen cells were collected on day 21 in the group without RSV challenge. (B) Endpoint titers of immunized antigen-binding IgG in murine sera on day 28 determined by ELISA. (C)-(D) Neutralization titers in the immunized murine sera on day 28 against RSV A2 (C) and RSV B9320 (D) strains. (E) IFNγ-secreting dots in mice spleens upon re-stimulation with indicated peptide pool (red and green indicated pool from monomer A and monomer B, respectively) assessed using ELISpot. (F)-(G) Equivalent RSV A2 (F) or B9320 (G) titers of lung and nasal tissues on 5 dpi determined by qRT-PCR. For panels (B) to (E), data were from two technical replicates of a single experiment. For panels (F) and (G), data were from three technical replicates of a single experiment. The dotted lines indicate the LLD. Any value less than the LLD was assigned a value of 1⁄2 the LLD. Bars represent mean ± SD. *P* values were analyzed with One-way ANOVA (ns indicated *P* > 0⋅05; * indicated *P* < 0⋅05; ** indicated *P* < 0⋅01, *** indicated *P* < 0⋅001, **** indicated *P* < 0⋅0001).

After the challenge with 1 × 10^6^ TCID_50_ RSV A2 or B9320 stain, high levels of virus RNA were detected in all tissues of LNP control mice. In contrast, the vaccinated groups exhibited approximately 2-3 orders of magnitudes lower virus titers in lung tissues (Figure 4F, G). Vaccination also resulted in a 20-fold reduction in viral load in nasal tissues. No significant differences were observed between the two vaccination groups in terms of viral loads in both tissues. Additionally, we also monitored and compared viral loads in lung and nasal tissues between the scDimer AB-based mRNA and LNP control groups. Using qRT-PCR-based viral load determination, we consistently observed lower viral loads in the scDimer AB-based mRNA immunization group compared to the LNP group at every detection time point (Figure S8). These data suggested that scDimers-mRNA were also effective in inhibiting RSV replication and facilitating virus clearance.

### Intramuscular scDimer AB mRNA-LNP followed by intranasal scDimer AB protein provided effective protection against viral challenge in mice

The COVID-19 outbreak highlighted the importance of mucosal immunity and the development of mucosal vaccines.^26^ We combined intramuscular and intranasal immunization to assess mucosal immune activation. Using the scDimer AB construct, mice received LNP-encoded vaccine with booster via intramuscular administration of mRNA vaccine (AB-IM) or intranasal administration of purified protein (AB-IN). Negative controls received empty LNP, while a separate group received intramuscular FI-RSV to induce vaccine-enhanced disease (VED) symptoms (Figure 5A).

**Figure 5.**
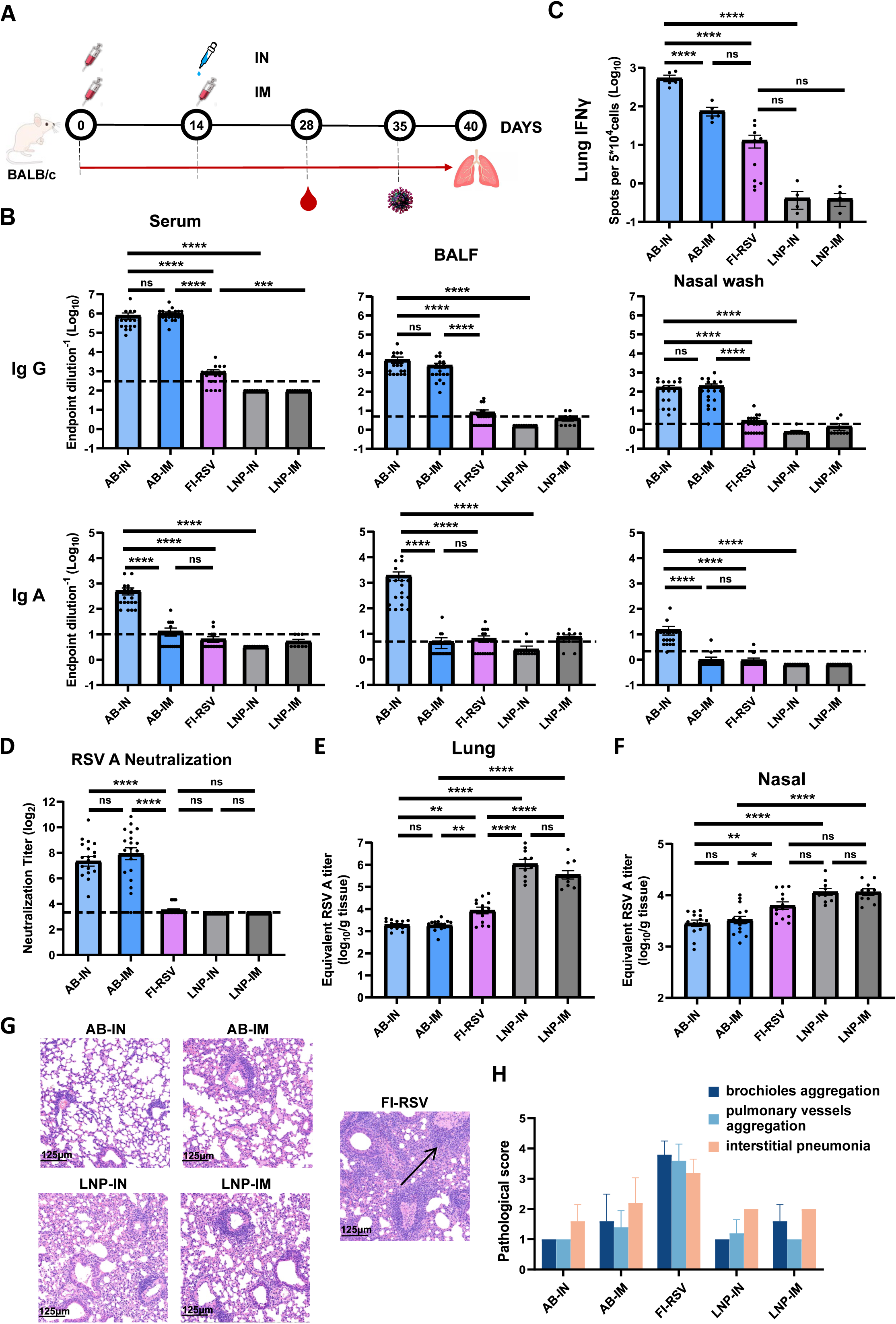
Immunization with intramuscular and intranasal delivery of scDimers against lung damage caused by RSV infection. (A) Procedure for mice immunization and sample collections. After the initial intramuscular administration of mRNA-LNP encoding scDimer AB, groups of BALB/c mice received a booster dose via either intramuscular injection (IM group) or intranasal delivery (IN group) of purified scDimer AB subunit protein vaccine. Two doses of intramuscular FI-RSV were administered at a 14-day interval as vaccine-enhanced disease (VED) controls. The immunized mice were then challenged with intranasal RSV subtype A2 on 35 days after the first dose (n = 15～20). Lung and nasal tissues were harvested at 5 dpi. (B) Endpoint titers of IgA and IgG in murine nasal washes, BALF, and serum were determined by ELISA on 28 days after the first dose. (C) IFNγ-secreting spots in lungs were assessed on day 28 upon re-stimulation with the AB peptide pool (a mixture of monomer A and monomer B) using the ELISpot assay. (D) Neutralization titers in the murine sera against RSV A2 strains on day 28. (E)-(F) Equivalent RSV A2 titers of lung (E) and nasal (F) tissues on 5 dpi. (G)-(H) Histological pathology analyses of lung sections from mice challenged with RSV A2 strain. Pathological scores (G) for immunocyte aggregation around bronchioles and pulmonary vessels, as well as interstitial pneumonia, were evaluated based on H&E-stained sections (H). For (B) to (D), data were from two technical replicates of a single experiment. For (E) and (F), data were from three technical replicates of a single experiment. The dotted lines indicate the LLD. Any value less than the LLD was assigned a value of 1⁄2 the LLD. Bars represent mean ± SD. *P* values were analyzed with One-way ANOVA (ns indicated *P* > 0⋅05; * indicated *P* < 0⋅05; ** indicated *P* < 0⋅01, *** indicated *P* < 0⋅001, **** indicated *P* < 0⋅0001).

IgA is the dominant antibody at many mucosal sites following mucosal vaccination. We evaluated RSV-specific binding IgA and IgG titers in nasal washes, bronchoalveolar lavage fluid (BALF), and serum (Figure 5B). Mice in AB-IN group developed higher anti-AB IgA titers in all three tissues compared to the other immunization groups, whereas IgG titers inAB-IN and AB-IM groups are comparable. Notably, antibodies in both AB-IN and AB-IM groups predominantly bound to the scDimer protein that maintains the pre-F conformation, with minimum binding to the post-F form. In contrast, FI-RSV primarily elicited post-F-specific antibodies (Figure S9A). To compare the differences in T cell immunity between AB-IN and AB-IM immune strategies in both local respiratory mucosal and systemic immunity, we quantified IFNγ-secreting cells isolated from lung and spleen tissues using ELISpot assay after 18 hours peptide stimulation. The results indicated that compared with AB-IM group, AB-IN significantly enhanced T-cell immunity in the respiratory mucosa of lung. However, systemic IFNγ levels (indicated by the IFNγ-screting spleen cells) of AB-IN group were markedly lower than those observed in AB-IM (Figure 5C, S9B).

Regarding the humoral immunity, high levels of neutralization antibodies against both subtype A and B were also observed in AB-IM and AB-IN groups. However, no detectable neutralization was observed in FI-RSV group, which might be attributed to the sensitivity limitation of the classic CPE assay (Figure 5D, Figure S9C). To precisely evaluate the neutralizing titers of FI-RSV group, we further conducted a highly sensitive fluorescence-based assay using GFP-tagged RSV A long strain, and the neutralizing titers were determined as 20-156 (Figure S9D).

When challenged with the RSV A2 strain, the immunized groups exhibited a significant reduction in viral loads within lung tissues, with ∼569-fold decrease in AB-IN group compared to LNP-IN group, and ∼195-fold decrease in AB-IM group compared to LNP-IM group (Figure 5E). In addition, both AB–IM and AB-IN group exhibited markedly lower viral loads in nasal tissues (Figure 5F).

Based on current knowledge, FI-RSV immunization induced imbalanced T cell responses, where mice vaccinated with FI-RSV demonstrated enhanced lung recruitment of conventional CD4^+^ T cells, and with higher Th2 but lower Th1 stimulation, which might partially contribute to VED.^27^ Following the RSV A2 challenge, both AB-IM and AB-IN groups exhibited diminished CD4^+^ T cell alongside increased stimulated CD8^+^ T cells (Figure S9E-G) in the lungs compared with FI-RSV group. These findings indicate a shift toward lower Th2 and higher Th1 responses, suggesting a decreased risk for VED. Consistent with these observations, lungs from FI-RSV-immunized mice displayed obvious immunocyte aggregation around bronchioles and pulmonary vessels (as indicated by black arrows), typical of VED (Figure 5G, H).^27^ Conversely, no apparent phenomenon was observed in both AB-IM and AB-IN groups, similar to LNP control group (Figure 5G). Interstitial pneumonia scores were also evaluated based on H&E-stained sections. FI-RSV received the highest scores. Both AB-IN and LNP-IN groups received comparable scores, likely lower than AB-IM and LNP-IM groups. In summary, both AB-IM and AB-IN vaccination strategies appear to have avoided the occurrence of VED, suggesting their potential as safe and effective candidates for RSV vaccines.

## DISCUSSION

Vaccination is a major prophylactic strategy to combat infectious diseases. The challenges associated with developing an inactivated RSV virus vaccine, along with the risk of VED have driven the pursuit of innovative solutions,^28^ particularly through structural vaccinology. The success of the pre-F stabilized subunit vaccines in the elderly highlights the effectiveness of this approach. However, various studies have reported the instability of soluble DS-Cav1,^14,15^ with pre-F-specific MAbs binding decreased by 50 % after approximately 40 days of storage at 4 ℃.^14^ To enhance the antigen’s stability, we engineered the F-based subunit vaccines derived from the RSV A pre-F state. Our primary goal was to retain the most potent neutralizing epitopes Ø and V while stabilizing the prefusion conformation of the antigen. To achieve these goals, we removed the C-terminal refolding region and two relatively weak epitopes I and IV, while preserving partial epitope III for stabilization. The resulting antigen still maintained the conformation of the pre-F (PDB: 5k6i) head region and avoided any allostery effects. Employing our tandem-antigen-repeat strategy, previously proven successful in vaccines against SARS-CoV-2 and MPXV,^23,24^ we fused the two designed F subunits to create a distinct molecular architecture: a single polypeptide chain engineered to fold into a stable dimeric conformation (scDimer). This design fundamentally differs from conventional dimers formed through intermolecular association, as it maintains covalent linkage while preserving key structural features. These scDimers interacted with MAbs as expected and induced robust humoral and cellular immune responses. Remarkably, after 100 days of storage at 4 ℃ and 28 days at 37 ℃, the average residual association of the antigen with epitopes Ø, V, and II still exceeded 50 %, demonstrating its exceptional stability in the pre-F state. Concerns may arise regarding neo-epitopes, as residues typically buried within the trimer structure are accessible in our design. However, it is reported that the F trimers undergo “breathing” in solution, making those buried residues also accessible *in vivo*. The exact exposure of neo-epitopes and their potential impact on the Mab repertoire needs to be further validation in our future studies.

Given the high amino acid sequence homology of the F protein between RSV A and B,^16^ most antibodies targeting F protein exhibit cross-neutralization against both viral subtypes. However, previous study reported antibodies exclusively neutralizing RSV A, while some others exhibit potency solely against RSV B.^29^ Thus, we designed a bivalent antigen, scDimer AB. This approach aligns with the strategy of one licensed RSV vaccines (Abrysvo of Pfizer), which is also bivalent. Unlike Abrysvo, which uses a mixture of two F protein-based antigens, our scDimer AB fused pre-F-based proteins from both subtypes into a single construct. This design facilitates the simultaneous activation of antibodies and may ultimately result in a balanced immune response. The results showed that this design provides effective protection against infection from both RSV A and B, addressing the challenge of predicting the predominate subtype in any given year. However, it is important to note that in our experiments, DS-Cav1 also provides cross-protection against both RSV A and B subtypes. Since subtype-specific antibodies have been reported, further investigation is required to determine whether this cross-protection extends to humans.

In the early stages of RSV vaccine development, VED was a devastating outcome that significantly hindered progress for nearly 60 years.^28^ Even today, defining the precise factors contributing to VED remains a challenge. According to FDA reports, during the clinical trials of Moderna mRNA-1345 and mRNA-1365, severe lower respiratory tract infection (LRTI) occurred in 3 infants under two years old following immunization. Although the LRTI may be related to the underdeveloped immune system of infants and no causative link with VED has been established, such results highlight the importance to minimize VED risk in antigen design. In our construct, the focuse on epitopes enhances the likelihood of generating high potency neutralizing antibodies while reducing the potential risk of producing antibodies associated with adverse reactions, if any. Consistently, our results showed that compared with FI-RSV group, mice from both AB-IM and AB-IN groups exhibited significantly lower proportions of CD8^+^ T cell and higher proportion of CD4^+^ T cell. Moreover, AB-IM and AB-IN groups demonstrated reduced interstitial pneumonia scores, suggesting a lower risk of inducing VED.

The proportion of CD4^+^ T cells in the lungs of AB-IN was higher compared with AB-IM group. A similar phenomenon was also observed in previous study, where an intranasal SARS-CoV-2 spike booster induced significantly higher levels of lung resident antigen-specific CD4^+^ T cells, with enhanced proportion of CD4^+^ Th1 and Th17 cells, while CD4^+^ Th2 remained unchanged.^26^ The precise mechanisms governing the functional polarization and tissue-specific interactions of diverse subsets (The, Th2, Th17 and Treg) in VED pathogenesis remain incompletely defined. Therefore, the increased proportion of CD4^+^ T cells in the lungs of AB-IN group compared to AB-IM group may be attributed to an enhanced Th1 response rather than Th2, and does not necessarily relate to VED. Further investigations are needed to delineate the specific CD4^+^ T cell subsets recruited to the lung parenchyma following intranasal antigen administration and RSV infection, with a particular emphasis on elucidating the underlying mechanisms, including but not limited to chemokine-mediated trafficking, antigen-presenting cell interactions, and the transcriptional programs governing subset differentiation and tissue residency.

Mucosal immunity is highly effective strategy to prevent respiratory viral infections,^26^ as it significantly boosts IgA antibody levels in the nasal passages and lungs, thereby reducing both viral infection and transmission within populations. In our study, the immunization strategy of intramuscular injection of an mRNA vaccine followed by nasal instillation of unadjuvanted protein vaccine was more effective in enhancing IgA titers in the nasal washes and BALF compared to a two-dose intramuscular injection of the mRNA vaccine group. These findings highlight the potential impact of mucosal vaccine on preventing lower respiratory tract infections and limiting RSV transmission via airborne.

Our study has some limitations. First, we only used DS-Cav1 as the positive control and did not include licensed vaccines for comparison of immunogenicity with our antigens, which will be addressed in future studies. Second, the RSV vaccines currently used in clinics, particularly for the elderly, primarily served as a booster to pre-existing immunity from natural infection.^30^ Meanwhile, our experiments were conducted on naïve animals, which may not fully reflect the real-world application scenario. Third, evaluating VED in rodent models is challenging, and future evaluations could be performed in non-human primates. Therefore, while our study provides a conceptual proof of principle, further investigation is needed before clinical application.

In summary, we report a novel subunit design inhibiting pre-F transition to post-F, which is promising for industrial development due to its high yield and thermal stability. The scDimer-based RSV vaccines effectively protected rodents from RSV infections, highlighting their potential for clinical application. This study proposed an immunogen design strategy that preserves potent neutralizing epitopes while eliminating weaker ones, resulting in immunogen modules that effectively present robust immune epitopes. By fusing various dominant immunogen modules, this approach enables their synergistic presentation to the immune system, promoting a balanced activation of the host immune response and providing broad-spectrum protection. Our strategy provides a novel framework for developing next-generation vaccines through the construction and combination of stable, optimized antigen modules, which can be further applied to other pathogens including, but not limited to hMPV, influenza, etc.

## Supporting information

method details, supplemental figures

## ACKNOWLEDGMENTS

We thank the staff of beamline BL02U1 and BL19U1 at the Shanghai Synchrotron Radiation Facility for assistance during crystal data collection. We are grateful to Bingxue Zhou, Yuanyuan Chen, and Zhenwei Yang from the Institute of Biophysics, Chinese Academy of Sciences, for their technical support in SPR and Octet assay. We thank Lin An, He Su, and Zheng Fan from the Institute of Microbiology, Chinese Academy of Sciences, for the protein yield and thermal stability assay. We also thank the Chinese Center for Disease Control and Prevention, and Laboratory Animal Center, for experimental animal feeding, including Mei Liu, Zijian Cheng, and Mingxuan Lan.

This work was supported by the National Key R&D Program of China (2020YFA0509202 and 2021YFC2301400 to J.Q.) and the National Science Foundation of China (82225021 to Q.W, 92169208 to J.Q. and 32200119 to Q.W). Q.W. is supported by Young Scientists in Basic Research, Chinese Academy of Sciences (YSBR-010). Special Program of China National Tobacco Corporation (2024200CB0160, 2024600CB0070 and 110202102034).

## AUTHOR CONTRIBUTIONS

Q.W., G.F.G., and J.Q. developed the antigen concept and managed various aspects of this study; J.L. and W.G. express the proteins and finished the protein character assays; J.L., W.G. and J.Z. crystalized proteins; X.M. did the mRNA in vitro transcription and lipid-nanoparticle encapsulation; J.L., X.M., Z.X., W.G., Y.Z., Y.M., J.Z., Q.W., S.L., J.C., Y.G., Q.W., and X.L. performed the animal experiments; J.L., X.M., W.G., J.C., Y.L., Y.Z. and S.L. cultured the HEp-2 cells and amplified RSV; J.L., X.M., Q.W., Y.G., and M.J. drafted the manuscript; J.L., X.M., Z.X., W.G., R.P., P.D., Q.W., G.F.G., and J.Q reviewed and edited the manuscript.

## SUPPLEMENTAL INFORMATION

Method details

Figures S1-S9.

