## Supplementary material for "Rational design of respiratory syncytial virus dimeric F-subunit vaccines in protein and mRNA forms": method details, supplemental figures

#### Cells

Expi293F cells were cultured at 37 °C in SMM 293-TII Expression Medium with 5 % $CO_2$ in a shaking incubator (140 rpm). HEK293T cells (ATCC) and HEp-2 cells (ATCC) were cultured at 37 °C in Dulbecco's Modified Eagle medium (DMEM) supplemented 10 % with fetal bovine serum (FBS) with 5 % $CO_2$.

#### Virus

RSV A2, an A subtype of RSV, was kindly provided by Professor Jianwei Wang (Key Laboratory of Respiratory Disease Pathogenomics, Chinese Academy of Medical Sciences & Peking Union Medical College, Beijing, China), which was purchased from ATCC (VR-1540). RSV B9320, a B subtype of RSV, was kindly provided by Professor Wenbo Xu (National Institute for Viral Disease Control and Prevention, Chinese Center for Disease Control and Prevention, Beijing, China), which was purchased from ATCC (VR-955). All RSV strains were propagated in HEp-2 cells in DMED supplemented with 2 % FBS and preserved in liquid nitrogen before used for antiviral activity assays and combination studies.

#### Mice and Cotton Rats

Specific pathogen-free (SPF) female BALB/c mice, aged 6 to 8 weeks, were purchased from Beijing Vital River Laboratory Animal Technology Co., Ltd. Similarly, SPF female Cotton Rats of the same age range were obtained from SPF (Beijing) Biotechnology Co., Ltd. All animals were allowed free access to water and standard chow diet, and were subjected to a 12-hour light-dark cycle (temperature: 20-25 °C, humidity: 40-70 %). The animals used in this study were in good health and had not undergone prior experimental procedures. They were housed under SPF conditions at the laboratory animal facilities of the Institute of Microbiology, Chinese Academy of Sciences (IMCAS), and the Chinese Center for Disease Control and Prevention (China CDC), Laboratory Animal Center. Mice were housed with 5 or 6 companions per cage, while cotton rats were housed with 2 companions per cage. Challenge experiments with RSV were conducted in an animal biosafety level 2 (ABSL2) facility at China CDC, Laboratory Animal Center. All animal experiments were approved by the Committee on the Ethics of Animal Experiments of IMCAS (Application Number: SQIMCAS2022091) and were conducted in accordance with the recommendations outlined in the Guide for the Care and Use of Laboratory Animals of the IMCAS Ethics Committee.

#### Protein Expression and Purification

For each construct, the signal peptide sequence of the RSV F protein (protein residues 1-25) was added to the protein's N terminus for secretion, and a hexa-His tag was attached at the C terminus for convenient purification. These constructs were codon-optimized for mammalian cell expression and synthesized by Azenta Life Sciences and GenScript, China, respectively. They were then individually cloned into the pCAGGS vector and transiently transfected into Expi293F cells. After 5 days, the supernatant was collected, and the soluble protein was purified using Ni affinity chromatography with a HisTrap™ HP 5 mL column (GE Healthcare). The bound protein was detached from the column using a buffer containing 0·3 mM imidazole, 20 mM Tris-HCl (pH 8.0), and 150 mM NaCl. Further purification was carried out by size-exclusion chromatography (SEC) using either a HiLoad™ 16/600 Superdex™ 200 pg or a Superdex™ 200 Increase 10/300 GL column (GE Healthcare) in a buffer composed of 20 mM Tris-HCl (pH 8·0) and 150 mM NaCl. Finally, SDS-PAGE was performed to analyze the size and purity of the eluted peak proteins.

Each plasmid of RSV F epitope monoclonal antibodies (MAbs), including 4D7, Palivizumab, 101F, CR9501 and AM22, were transiently transfected into Expi293F cells with pCAGGS vector containing immunoglobulin heavy chain and light chain sequences respectively. After 5 days, the supernatant was collected and purified using HiTrap™ Protein A HP column (Cytiva). The MAbs were eluted from the Protein A column using 0·1 M glycine, pH 3·0, and neutralized by adding 0·25 times the volume of 1 M Tris (pH 9·0) to each elution tube. After concentrating the protein-buffer mixture, further purification was conducted by size-exclusion chromatography (SEC) using PBS. To generate antibody-binding fragment (Fab), purified 4D7, Palivizumab, 101F, CR9501, and AM22 MAbs were digested with immobilized papain (Thermo Scientific) following the manufacturer's instructions. Fab fractions were then purified using a HiTrapTM Protein A HP column (Cytiva) with PBS. The plasmid encoding the

MAb of site III, MPE8 Fab, was transfected into Expi293F cells using a pCAGGS vector containing only the Fab fragment sequence with a hexa-His tag at the C terminus. After 5 days of suspension culture, the protein was purified using a HisTrap$^{TM}$ HP 5 mL column followed by size-exclusion chromatography using Superdex$^{TM}$ 200 Increase 10/300 GL in PBS.

#### Crystallization, Data Collection and Structure Determination

Before crystallization, RSV F monomer A and scDimer AA were deglycosylated by PNGase F[1] in room temperature for 3 hours. This hydrolase was expressed in E. coli BL21 and purified using a HisTrap$^{TM}$ HP 5 mL column followed by size-exclusion chromatography using HiLoad$^{TM}$ 16/600 Superdex$^{TM}$ 75 pg in PBS.

Crystals of the deglycosylated proteins were grown by sitting-drop method with 0·8 µL protein mixing with 0·8 µL reservoir solution at 18 °C. General crystallization screenings were performed with commercially available kits. Diffractable crystal of monomer A was obtained in the reservoir solution consisting of 0·1 M phosphate/citrate (pH 4·2), 40 % v/v PEG300, with protein concentration of 10 mg/mL. X-ray diffraction data were collected at Shanghai Synchrotron Radiation Facility (SSRF) BL02U1. Diffractable crystal of scDimer AA was obtained in the solution consisting of 1·5 M sodium chloride, 10 % v/v ethanol, with protein concentration of 10 mg/mL. X-ray diffraction data was collected at SSRF BL19U1. For data collection, the crystals were cryo-protected by briefly soaking in reservoir solution mixed with 20 % v/v glycerol before flash-cooling in liquid nitrogen. The datasets were processed using HKL2000 software.[2] The structures of monomer A and scDimer AA were determined by the molecular replacement method using Phaser with previously reported RSV pre-F structure (PDB: 5k6i).[3] The atomic models were completed using Coot[4] and refined with phenix.refine in Phenix.[5] The stereochemical qualities of the final models were assessed with MolProbity.[6] Data collection, processing, and refinement statistics are summarized in Table 1. All structural figures were generated using Pymol software (https://pymol.org).

#### Surface Plasmon Resonance (SPR) Assay

Monomer A and scDimer AA were biotinylated and transferred into PBST buffer (1·8 mM $KH_2PO_4$, 10 mM $Na_2HPO_4$ (pH 7·4), 137 mM NaCl, 2·7 mM KCl, and 0·05 % (v/v) Tween 20), then immobilized on an SA chip. The MAbs Fab fragments were serially diluted and then flowed over the chip in PBST buffer. Binding affinities were measured by BIAcore 8K (GE Healthcare) at 25 °C in single-cycle mode. Binding kinetics were analyzed with Biacore Insight software (GE Healthcare) using a 1:1 Langmuir binding model to calculate KD values. The graphics were prepared with GraphPad Prism 9.

#### Differential Scanning Fluorimetry (DSC)

To evaluate the physical stability of the prefusion conformation of the designed RSV F glycoproteins under thermal stress, the proteins' melting temperature (*T*m) was determined using differential scanning fluorimetry performed on a Prometheus NT.48 instrument (Nano Temper), a high-throughput method measuring 48 samples at the same time. The scDimer AA, diluted to 2 mg/mL in PBS buffer, was dispensed into standard capillary tubes in 10 μL aliquots. The scanning temperature range was set from 20 to 95 °C with a heating rate of 1 °C/min. Protein unfolding was monitored by detecting changes in tryptophan fluorescence at emission wavelengths of 330 nm and 350 nm, depending on the temperature gradient. These scans were performed as single reads, and *T*m values were calculated according to the manufacturer’s instructions. All samples were run in duplicate.

The *T*m values of scDimer AA and AB were also measured by the classic DSC method on nano-DSC (TA instruments) with the same detection concentration and scanning temperature range.

Graphics were prepared using GraphPad Prism 9.

#### Conformation Stability Studies

To quantitatively analyze the stability of key epitopes of designed RSV F glycoproteins, the Octet-based method was employed. Palivizumab, CR9501, and AM22 MAbs were coated on Protein A biosensors at a concentration of 5 μg/mL. The antigens were stored at 4 °C or 37 °C for the indicated period in a solution buffer containing 20 mM Tris-HCl (pH 8.0) and 150 mM NaCl. Antigens from different time points were utilized at a concentration of 20 nM for quantitative experiments conducted on the Octet

Red96 (ForteBio) at 30 °C for 360 s. Data analysis was executed using ForteBio Octet DataAnalysis 9.0.10. Graphics were prepared using GraphPad Prism 9.

#### Protein Yield Assay

The plasmids containing each construct were cloned into the pCAGGS vector and transiently transfected into HEK293T cells in 6-well plates. Following three days of static culture without FBS at 37 °C and 5 % $CO_2$, the supernatant from each well was collected. Protein concentration, including a C-terminal His-tag, was determined using the BCA kit (Beyotime) and diluted two-fold from 6 μg/mL to create six gradients. Protein samples were mixed with Sample Buffer (Bio-Techne) and other components as per the Jess™ ProteinSimple instructions according to the instruction of Jess™ ProteinSimple. Serial dilutions were detected alongside the supernatants using the same sample volume. Data were collected and analyzed using Compass v6.1.0 (Bio-Techne) (http://proteinsimple.com/compass/downloads/). A standard curve was constructed based on the area under the curve (AUC) and the concentration of the serially diluted protein, enabling determination of the supernatants' concentration for calculating the protein yield.

#### mRNA in vitro Transcription and 5'-Capping

mRNA was transcribed in vitro using T7 High yield RNA transcription Kit (Novoprotein) on linearized plasmids encoding scDimers. A 57 nucleotide-long poly(A) tail was transcribed into mRNA. UTP was replaced by 1-methylpseudourine-5'-triphosphate during in vitro transcription to generate nucleotide-modified mRNA. 5'-Capping was conducted using Cap 1 capping system (Novoprotein). The mRNA was purified by precipitation with LiCl at -20 °C overnight, followed by centrifugation at 18,800 × g for 20 minutes at 4 °C, and resuspension with RNase free water. Purified mRNA was verified using agarose gel and stored at -80 °C until use.

#### Lipid-nanoparticle Encapsulation of mRNA

Lipid-nanoparticle (LNP) encapsulation was performed using the Nanoassemblr Benchtop platform (Precision Nanosystems). Essentially, mRNA, diluted in an aqueous solution (pH = 4.0), was rapidly mixed with lipids, diluted in ethanol, within a microfluidics system, facilitating the encapsulation of mRNA by lipids through a self-assembly process. The LNPs utilized in this study were assembled using an ionizable cationic lipid, phosphatidylcholine, cholesterol, and PEG-lipid at a ratio of 50:10:38·5:1·5 (mol/mol). The mRNA to lipid ratio was approximately 0·05 (wt/wt). Quantification of LNP-encapsulated mRNAs was conducted using Quant-iT RiboGreen RNA Assay Kit (Thermo Fisher). LNPs were stored at 4 °C at an RNA concentration of approximately 0.5 mg/mL.

#### mRNA Transfection and Western Blot

HEK293T cells were transfected using TransIT-mRNA kit (Mirus Bio). Speicfically, 1 μg of mRNA was added to 100 μl serum-free Opti- MEM, 2 μl of TransIT-mRNA reagent, and 2 μl booster reagent. The mixture was incubated for 2 min before being added dropwise to $5 \times 10^5$ cells cultured in complete medium in 12 well plates. Supernatants were collected 12, 24, 36 and 48 hours post-transfection and stored at −20 °C until further use. For western blot, supernatant samples were combined with loading buffer without dithiothreitol, separated by 10% SDS-PAGE and transferred to PVDF membrane using a semi-dry apparatus (WIX Technology). Then, the membrane was blocked with 5% non-fat milk diluted in PBS-T buffer. Serum from mice immunized with scDimer-AB was used as the primary antibody for 1 hour, followed by incubation with goat anti-mouse IgG-HRP secondary antibody (Easybio) for 1 hour. Finally, the membrane was developed using Beyotime BeyoECL Plus (Beyotime Biotech).

#### Particle Size of LNPs

To determine the particle sizes of LNPs, they were diluted into deionized water at pH 7·4. The diluted LNPs were then added to a folded capillary zeta cell and loaded into the Zetasizer Pro (Malvern Panalytical). Particle size distribution was measured using dynamic light scattering.

#### FI-RSV Vaccine Preparation

RSV-infected HEp-2 cell culture supernatant was harvested from the flasks, then clarified by low-speed centrifugation (1,000 g, 10 min, 4 °C). Formalin was added at a final dilution of 1:4,000 and incubated at 37 °C for 3 days for inactivation. The formalin-inactivated culture medium was transferred to sterile tubes (Ultra-clear, Beckman) and subjected to ultracentrifugation (60,000 g) for 60 min. The resulting pellets were resuspended to 1/25 of the original volume in MEM without supplements. The vaccine was adsorbed to aluminum hydroxide (4 mg/mL) overnight at room temperature. The compounded material was pelleted by centrifugation (1,000 rpm for 10 min) and the pellet was resuspended in 1/4 of the original volume.

#### Animal Immunizations and Virus Challenges

In the protein-antigen study, all animals were immunized intramuscularly 3 times at weeks 0, 3, and 6 with varying doses of antigen mixed with either Alhydrogel® (InvivoGen) or AddaVax™ (InvivoGen) adjuvant. The frozen antigens, stored in PBS, were thawed on ice and mixed with an equal volume of AddaVax™ adjuvant. For the mixture of antigen and Alhydrogel® adjuvant, it was diluted with sterile PBS to achieve a 0·5 % concentration of alum. Blood samples were collected 2 days before the first and second immunizations, and 1 week before the third immunization. The serum was separated by centrifugation of whole blood and stored at -20 °C until use. All mice were euthanized at week 9, and spleen tissue was collected for ELISpot experiments. Cotton rats were intranasally administered 100 µL (50 µL/nare) of live RSV A2 or RSV B9320 strain ($1 \times 10^6$ $TCID_{50}$) at week 9. 5 days later, all rats were euthanized, and lung, trachea, and nasal tissue were collected for further assays.

In the mRNA-antigen study, all mice were immunized with two doses of mRNA vaccine (5 µg/mouse per dose) or empty LNP as a placebo control on day 0 and day 14. Blood samples were collected via retro-orbital blood collection at day 14 and by cardiac puncture at day 28. The blood samples were then centrifuged, and the serum in the supernatants was stored at -80 °C until use. For evaluating the cellular immunogenicity of mRNA vaccines, groups of mice were immunized with two doses of mRNA vaccine (5 µg/mouse per dose) or empty LNP as a placebo control on day 0 and day 14. Mice were sacrificed 7 days after the second immunization on day 21, and spleens were collected immediately afterward for the ELISpot experiment. Another batch of mice received intranasal administration of 100 µL (50 µL/nare) of live RSV A2 or RSV B9320 strain ($1 \times 10^6$ $TCID_{50}$) respectively on day 35. Five days later, all mice were euthanized, and lung and nasal tissues were collected for further assays.

All mice in the combined mRNA and protein study were initially immunized intramuscularly with mRNA vaccine (5 µg/mouse per dose), followed by a secondary immunization with either mRNA vaccine intramuscularly or protein vaccine intranasally on day 14. Subsequent steps were consistent with the mRNA vaccine study protocol.

#### RSV F or Post-F Binding ELISA

96-well flat-bottom plates (Corning, 3590) were coated with either pre-F or post-F[7] protein (3 μg/mL), incubated overnight at 4 °C. Plates were washed with PBST buffer 4 times between each step, and all incubations were performed at 37 °C for 1 hour. The plates were blocked with 100 μL of 5 % skimmed milk in PBS. Serum samples were diluted 1:50 and underwent a series of four-fold dilutions in 96-well round-bottom plates (Corning, 3799). Then, the serial dilution was added to protein-coated plates. After incubation and washing, mouse-sample-plates were incubated with 100 μL of goat anti-mouse IgG-HRP antibody (1:5,000 dilution, Easybio, BE0102-100). Similarly, cotton rat sample plates were incubated with 100 μL of chicken anti-cotton rat IgG antibody (1:5,000 dilution, ICL, CCOT-25A), followed by incubation with 100 μL of goat anti-chicken IgG-HRP (1:5,000 dilution, Bioss, bs-0310G-HRP). The plates were subsequently developed with 3,3',5,5'-tetramethylbenzidine (TMB) substrate (Beyotime, P0209). The reactions were stopped with 100 μL 2 M hydrochloric acid, and the absorbance was measured at 450 nm using a microplate reader (ThermoFisher). The endpoint titers were defined as the highest reciprocal dilution of serum to give an absorbance greater than 2.1-fold of the background values. Antibody titers below the limit of detection were determined as half the limit of detection.

All samples were run in duplicate, and an internal reference control was included for accuracy.

#### Enzyme-linked Immunospot (ELISpot) Assay

Mouse spleen and lung tissues were collected and ground to separate single cells. The cells were stimulated with a peptide pool (10 μg/mL individual peptide) designed according to the antigen sequence, consisting of 13-17-mers overlapped by 11 amino acids (https://www.hiv.lanl.gov/content/sequence/PEPTGEN/peptgen.html). ELISpot assays were conducted using mouse IFNγ, TNFα, IL-2, and IL-4 ELISpot kits from MABTECH, following the manufacturer's protocol. Spot counts were determined using an automatic ELISpot reader from CTL and images analyzed with ImmunoSpot 7·0.

#### RSV Neutralization Assay

A cytopathogenic effect (CPE) assay using HEp-2 cells was conducted to assess neutralizing antibody activities induced by antigens in animals against RSV infection. Briefly, equal volumes of serially diluted serum and 200 $TCID_{50}$ of RSV subtype strain (RSV A2 strain or RSV B9320 strain) were mixed and incubated for one hour at 37 °C. The mixture of serum and virus was then added to 20,000 HEp-2 cells per well in 96-well flat-bottom plates, followed by incubation at 37 °C with 5 % $CO_2$ for 4 days. CPE was recorded to determine the antibody neutralizing titer.

RSV-EGFP neutralization assay has higher sensitivity than the method of CPE, the construction of RSV-EGFP was performed as previously described. The neutralization assay also using HEp-2 cells, the serially diluted serum was incubated with the equal volume 200 PFU virus for one hour at 37 °C. The mixture of serum and virus was then added to 20,000 HEp-2 cells per well in 96-well flat-bottom plates, after 3 days of incubation at 37 °C under 5% $CO_2$. The fluorescence signals from RSV-EGFP were observed under CQ1 confocal image cytometer (Yokogawa).

All samples were run in duplicate, and an internal reference control was included.

#### Virus Loading Assay

Standard curves were generated using known titer virus N gene amplification. Lung, trachea, and nasal tissues from cotton rats were weighed and homogenized. A portion of the homogenate was used to extract virus RNA using a MagaBio plus virus RNA purification Kit (Bioer, BSC58S2DB) on a GenePure Pro 32E (Bioer), following the manufacturer's instructions. RSV A or RSV B specific quantitative reverse transcription-PCR (qRT-PCR) assays were conducted using a One Step PrimeScript™ RT-PCR Kit (TaKaRa, RR064B) on a QuantiStudio™ 5 System (ThermoFisher), following the manufacturer's protocol. Two sets of primers and probes were utilized to detect a region of the N gene of the viral genome,[8] respectively, with the following sequences:

RSVA-F, AGATCAACTTCTGTCATCCAGCCA;

RSVA-R, ATTGATACTTCCTAATTATGATGTGC;

RSVA-probe, 6-FAM-CACCATCCAACGGAGCACAGGAGAT-TAMRA-N;

RSVB-F, AAGATGCAAATCATAAATTCACAGGA;

RSVB-R, CACTATAAAGATACTTAAAGATGCTGGATATCA;

RSVB-probe, 6-FAM-AGGTATGTTATATGCTATGTCCAGGTTAGGAAGGGAA-TAMRA-N.

Another portion of the homogenate was clarified by centrifugation at 4,000 rpm for 10 minutes, and the virus titer in the supernatant was determined by CPE assay with HEp-2 cells. Supernatant samples were serially diluted (100 μL) and added to 20,000 HEp-2 cells per well in 96-well flat-bottom plates, followed by incubation at 37 °C with 5 % $CO_2$ for 1 hour. The supernatant was then replaced with 100 μL of fresh DMEM supplemented with 2 % FBS, and the cells were further incubated for 4 days at 37 °C with 5 % $CO_2$. CPE was observed to determine the virus neutralizing titer.

All samples were run in duplicate, and an internal reference control was included.

#### Flow Cytometry

To quantify T cells in lung tissue, single-lung-cell suspensions were initially blocked with recombinant CD16/32 to prevent nonspecific Fc receptor binding. Subsequently, cells were surface-stained with monoclonal antibodies specific to CD3-PE-Cy7, CD4-APC-Cy7, CD8α-PerCP/Cy5.5, and Zombie UV viability staining solution (BioLegend) at 4 °C for 30 minutes. Flow cytometric analysis was performed using a BD LSRFortessa flow cytometer (BD Biosciences) equipped with a high-throughput system. Data analysis was conducted using FlowJo 10.0 software.

#### Histology

The lungs of mice were excised and fixed in 4 % neutral buffered formalin for 48 hours at room temperature, followed by embedding in paraffin. Upon slicing, the tissues were stained with hematoxylin and eosin (H&E). Pathological changes were assessed and scored on a severity scale ranging from 1-4. Scores for inflammatory cell aggregation and interstitial pneumonia were as follows: 1 indicated normal naive parameters, 2 indicated slight and occasional cell aggregation, 3 indicated moderate cell infiltration, and 4 indicated moderate to severe and multifocal cell aggregation around bronchioles, pulmonary vessels, or air spaces of lung sections.

### Supplemental Figures:

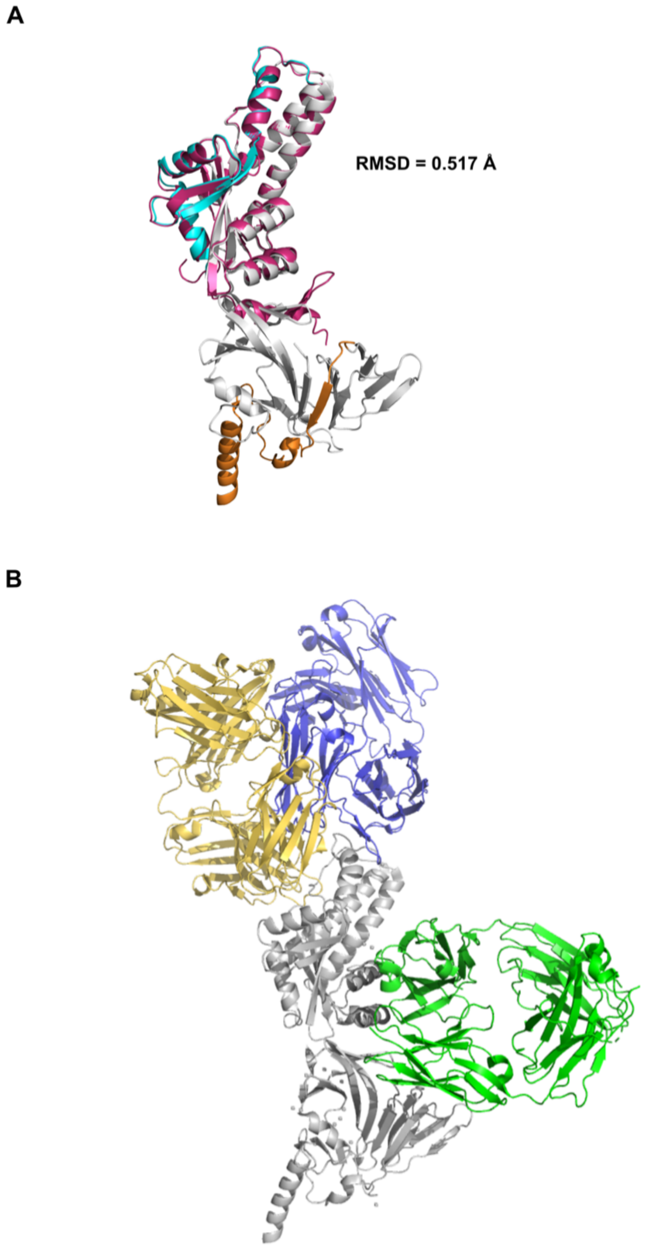


**Figure S1. Crystal structure align.**

(A) Crystal structure alignment of monomer A (pink) and F protein at prefusion state (gray, PDB: 5k6i). Refolding regions are colored in cyan and orange.

(B) Crystal structure alignment of F protein at prefusion state (PDB: 5k6i) and Fab segments of Ø, V and II epitope- antibodies. Am22 Fab (epitope Ø), CR9501 Fab (epitope V), Motavizumab (epitope II) are colored in blue, yellow and green, respectively.

A

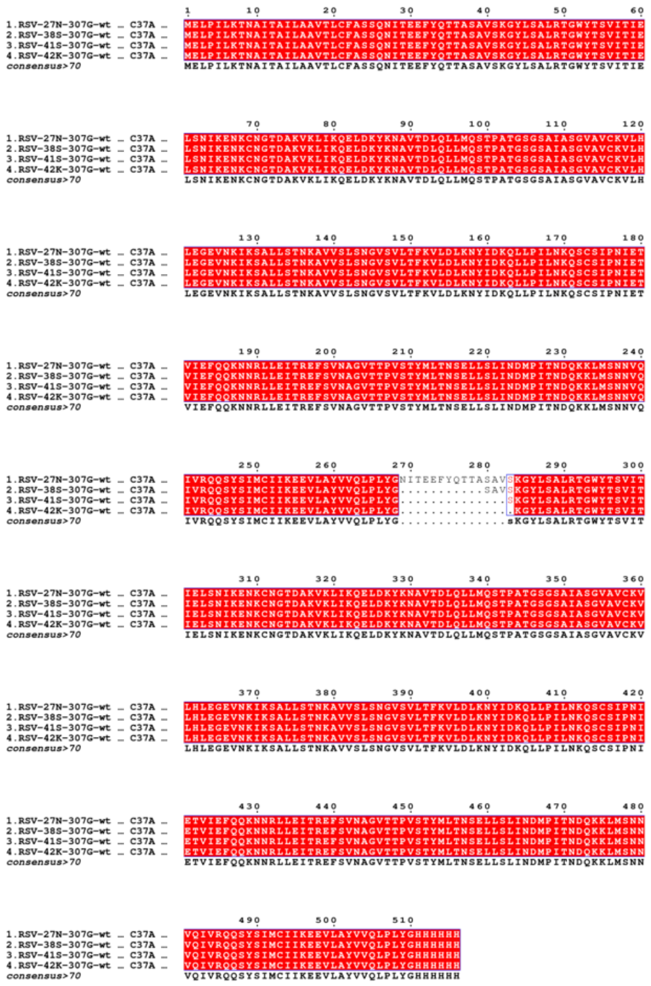

B

| Sample ID | Yield (mg/L) | *T*m (℃) |
|---|---|---|
| 1 | 65.55 ± 9.04 | 79.19 ± 0.41 |
| 2 | 6.02 ± 0.89 | 77.92 ± 0.19 |
| 3 | 6.30 ± 0.95 | 76.53 ± 0.03 |
| 4 | 6.82 ± 0.37 | 77.04 ± 0.04 |

**Figure S2. Characteristics of 4 scDimer designs**.
(A) Sequence alignment of four scDimer designs. Identical residues are highlighted in white on a red background, and residues in red on a white background indicate a similarity score >0·7, considering physio-chemical properties. The alignment was performed by T-COFFEE and visualized by ESPript 3·0.
(B) Yield and *T*m of scDimer designs. *T*m were determined using NT.48, a high-throughput method. Representative data of three replicates were shown.

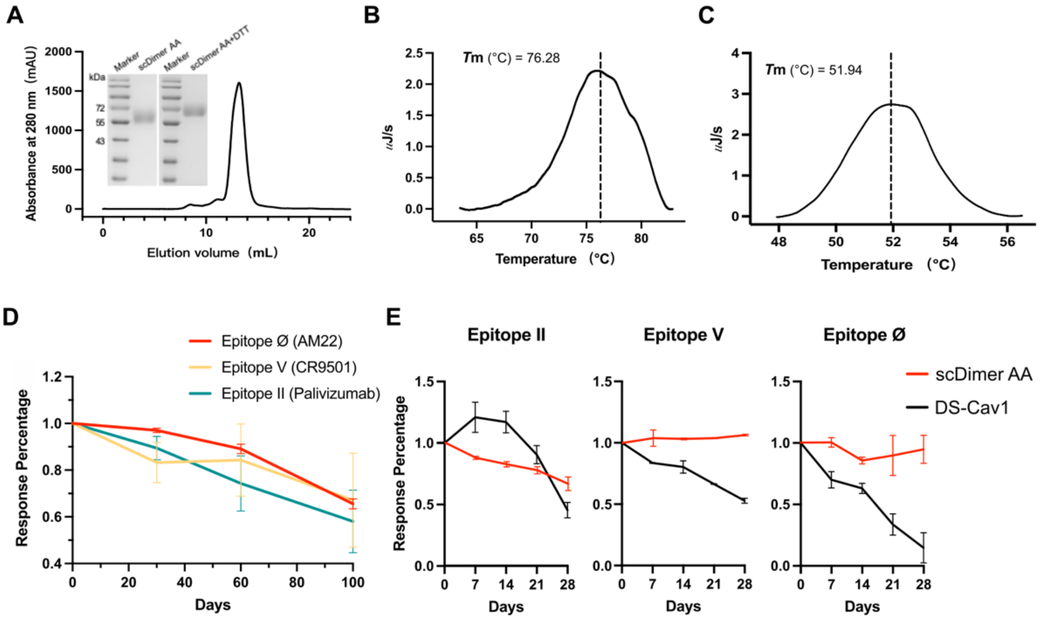


**Figure S3. Characterization of scDimer AA and DS-Cav1 Protein.**
(A) Size exclusion chromatography SDS-PAGE assay of scDimer AA.
(B) Thermal unfolding profiles of scDimer AA were determined using DSC. The dot line and values in graph represent obtained *T*m (maxima) of the main peak using a non-two-state model with two.
(C) Thermal unfolding profiles of DS-Cav1 were determined using DSC. The dot line and values in graph represent obtained *T*m (maxima) of the main peak using a non-two-state model with two.
(D) Stability evaluation of scDimer AA at 4 °C storage.
(E) Stability evaluation of scDimer AA and DS-Cav1 at 37 °C storage.
(D)-(E) Using the binding of pre-F-specific MAbs as the indicators, the proteins were bound to key epitope MAbs after storage at 4 °C or 37 °C for the indicated days or weeks. Bars and symbols represent the average of three measurements, while lines depict the range of values. The pre-F harvest (day 0) was normalized to 1.

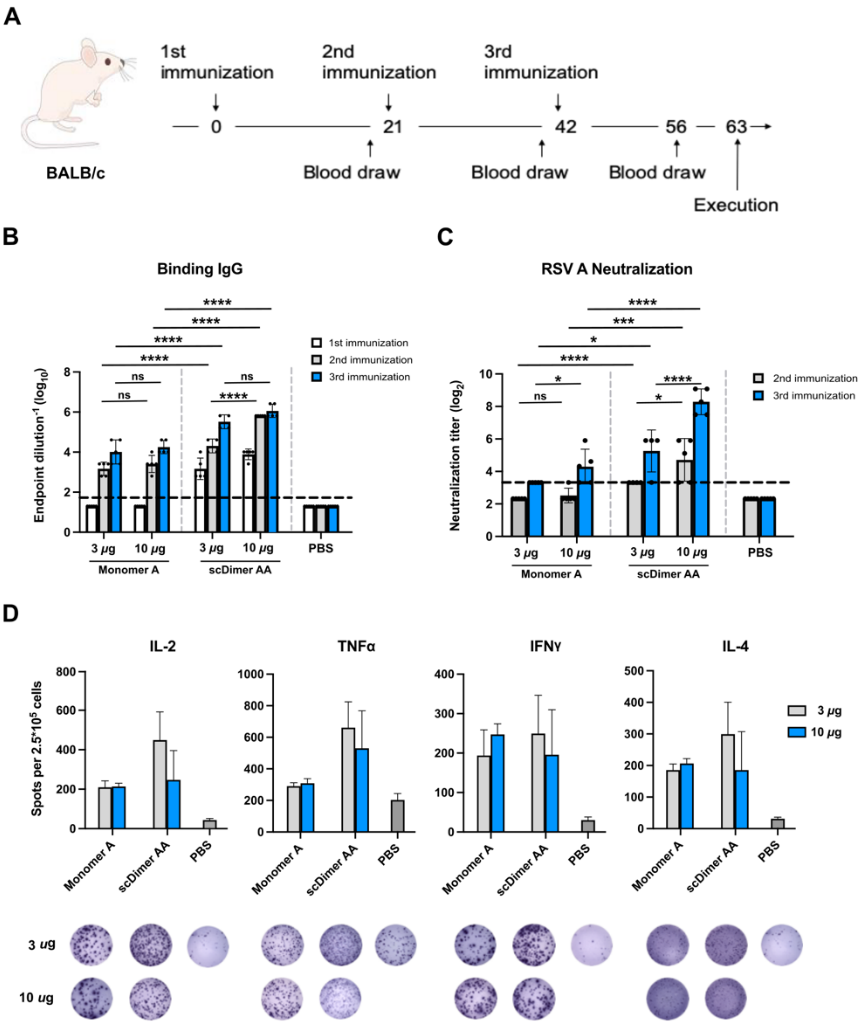


**Figure S4. Immunogenicity of Monomer A and scDimer AA in Mice.**

(A) Procedure for mice immunization, execution and sample collections. Groups of 6 to 8-week-old female BALB/c mice (n=4~5) were intramuscularly immunized with 3 or 10 μg of monomer A or scDimer AA on day 0, 21 and 42 with Alhydrogel® adjuvant. Serum samples were collected on day 19, 40 and 56. Splenocytes were collected on day 63.

(B) Endpoint titers of antigen-binding IgG in murine sera. Related to Figure 1C.

(C) Neutralization titers in the murine sera against RSV A2. Related to Figure 1D.

(D) Cellular immunity was evaluated. Groups of 6 to 8-week-old female BALB/c mice (n=4~5) were immunized with two doses of antigen adjuvanted with AddaVax™. Mouse IFNγ, TNFα, IL-2, and IL-4 ELISpot assays were performed to indicate monomer antigen peptide pool re-stimulation.

Data were collected from two technical replicates of a single experiment. Dotted lines indicate the lower limit of detection (LLD). Any value less than the LLD was assigned a value of 1⁄2 the LLD. Bars represent mean ± standard deviation (SD). *P* values were analyzed with Two-way ANOVA (ns indicated $P > 0{\cdot}05$; * indicated $P < 0{\cdot}05$; ** indicated $P < 0{\cdot}01$, *** indicated $P < 0{\cdot}001$, **** indicated $P < 0{\cdot}0001$).

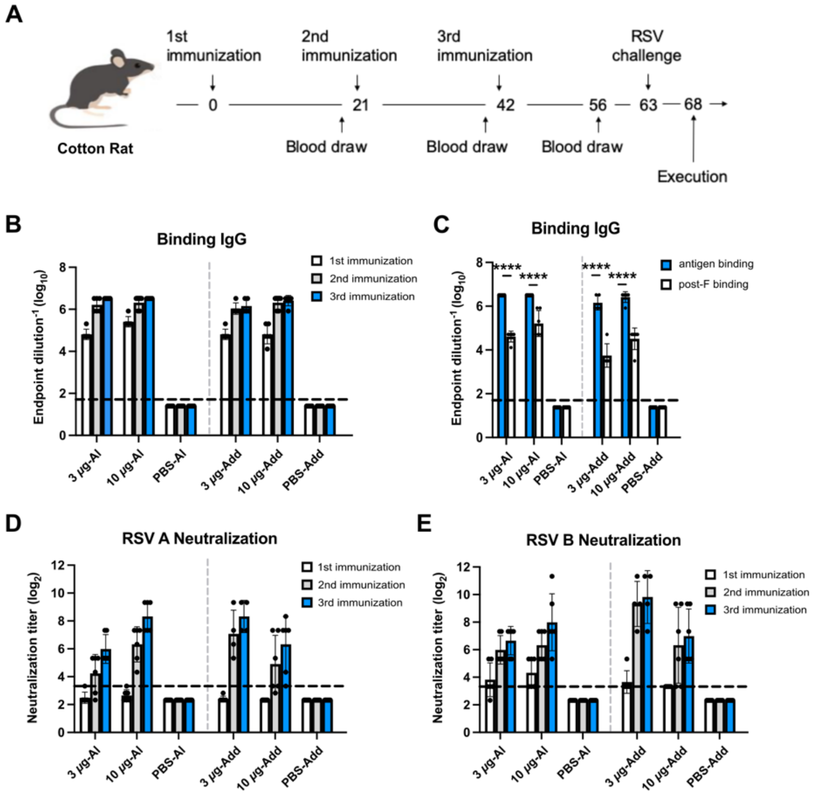


**Figure S5. Immunogenicity of scDimer AA in Cotton Rats.**

(A) Procedure for mice immunization, challenge and sample collection. Groups of 6 to 8-week-old female cotton rats (n=4~6) were intramuscularly immunized with 3 or 10 μg of scDimer AA on day 0, 21 and 42 with Alhydrogel® or AddaVax™. Serum samples were collected on day 19, 40 and 56. The immunized rats were then challenged with intranasal administration of $1\times10^5$ $TCID_{50}$ of RSV subtype A2 on day 65. Lung, trachea and nasal tissues were harvested 5 days post infection (dpi).

(B) Endpoint titers of antigen-binding IgG in sera of cotton rats.

(C) Endpoint titers of scDimer AA (blue column) or post-F (white column)-binding IgG in sera.

(D)-(E) Neutralization titers in the murine sera against RSV A2 (D) and RSV B9320 (E) strains. Related to Figure 2A, B.

For (B) to (E), data were collected from two technical replicates of a singlne experiment. Dotted lines indicate the LLD. Any value less than the LLD was assigned a value of 1/2 the LLD. Bars represent mean ± SD. *P* values were analyzed with Two-way ANOVA (ns indicated *P* > 0·05; * indicated *P* < 0·05; ** indicated *P* < 0·01, *** indicated *P* < 0·001, **** indicated *P* < 0·0001).

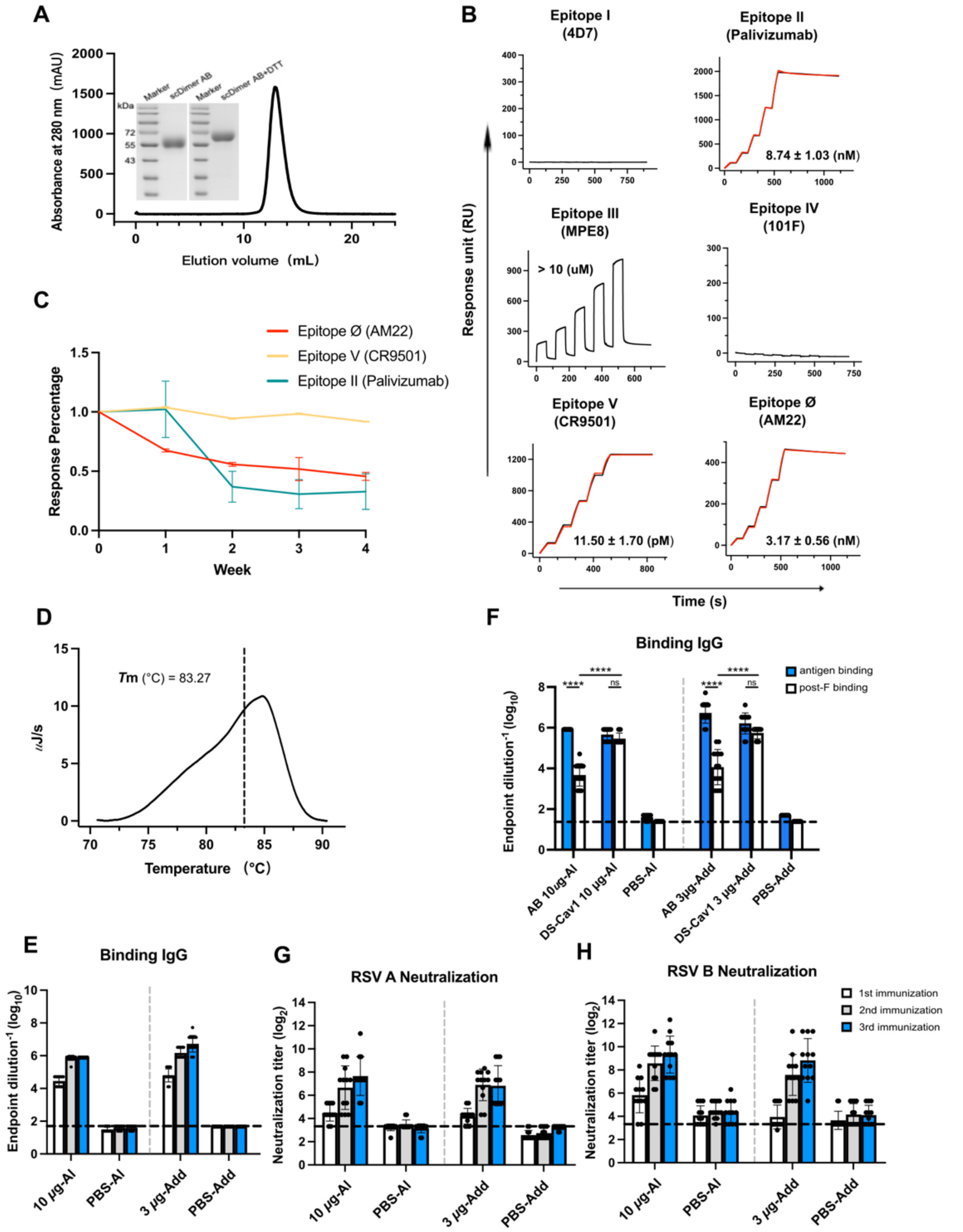


**Figure S6. Characterization of scDimer AB protein and its Immunogenicity Evaluations in Cotton Rats.**

(A) Size exclusion chromatography and SDS-PAGE of scDimer AB.

(B) Binding affinities of scDimer AB with MAbs targeting six epitopes. Actual and fitted curves were represented by red and black lines, respectively. $K_D$ values were listed as mean ± SD of three experiments.

(C) Stability evaluation of scDimer AA at 37 °C. Using the binding of pre-F-specific MAbs as the indicators, the protein was bound to key epitope MAbs after storage at 37 °C for the indicated days or weeks. Bars and symbols represent the average of three measurements, while lines depict the range of values. The pre-F harvest (day 0) was normalized to 1.

(D) Thermal unfolding profiles of scDimer AB (black) and fit (red) were determined using DSC. The dot line and values in graph represent obtained $T$m (maxima) of the main peak using a non-two-state model with two.

(E) Endpoint titers of antigen-binding IgG in sera of cotton rats.

(F) Endpoint titers of scDimer AB (blue) and post-F (white)-binding IgG in sera of cotton rats.

(G)-(H) Neutralization titers in the murine sera against RSV A2 (G) and RSV B9320 (H) strains. Related to Figure 3B, C.

For (E) to (H), data were collected from two technical replicates of a single experiment. Dotted lines indicate the LLD. Any value less than the LLD was assigned a value of 1⁄2 the LLD. Bars represent mean ± SD. $P$ values were analyzed with Two-way ANOVA (ns indicated $P > 0{\cdot}05$; * indicated $P < 0{\cdot}05$; ** indicated $P < 0{\cdot}01$, *** indicated $P < 0{\cdot}001$, **** indicated $P < 0{\cdot}0001$).

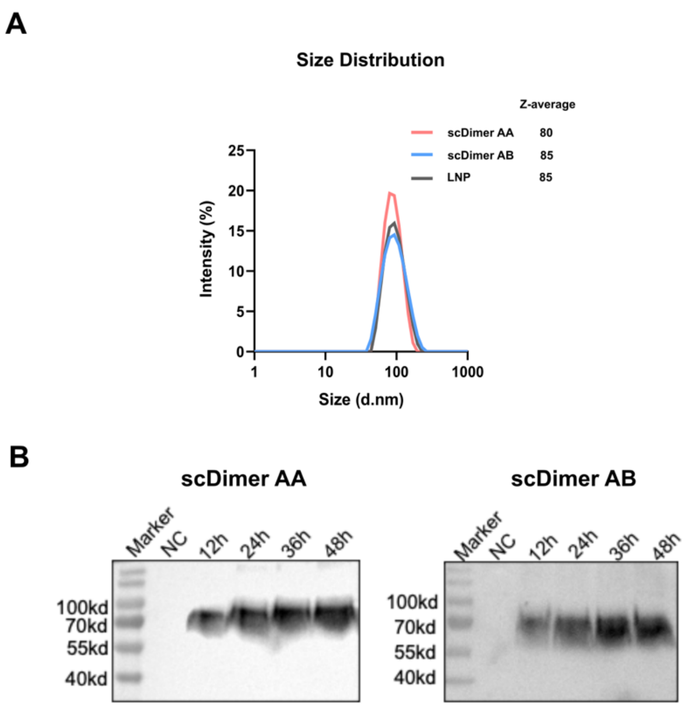


**Figure S7. Characterizations of scDimers LNP-particles and *in vitro* expression of the two mRNAs encoding scDimers.**

(A) Particle size distributions of LNPs characterized by dynamic light scattering (DLS). Numbers in legend indicate Z-average.

(B) scDimers mRNA was expressed in HEK293T cells. The supernatant was collected at 12, 24 36, and 48 hours and analyzed by western blotting. NC: negative control.

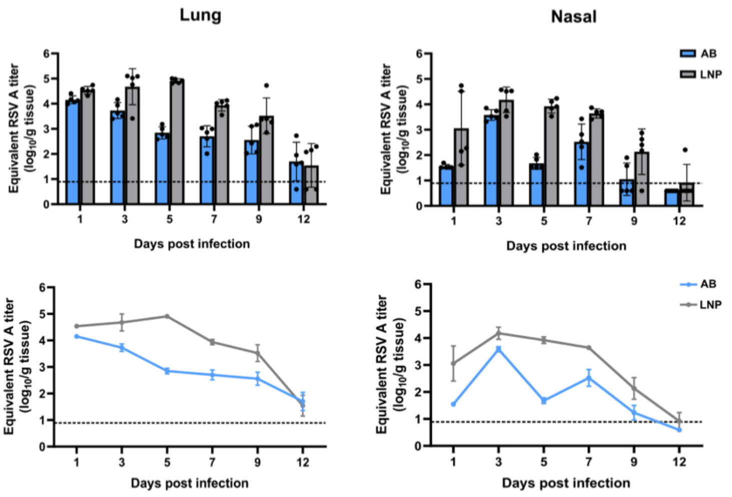


**Figure S8. Comparison of the viral clearance between the scDimer AB-based mRNA and LNP Control Groups.**

According to the immunization procedure depicted in Figure 4A, mice were euthanized at 1, 3, 5, 7, 9 and 12 days post-infection (dpi). Subsequently, the viral loads in homogenized lung and nasal tissues were determined by qRT-PCR. Groups of female BALB/c mice aged 6 to 8 weeks (n=5) were used at each time point. Data were obtained from three technical replicates of a single experiment. Dotted lines indicate LLD. Any value less than the LLD was assigned a value of 1⁄2 the LLD. Bars represent mean ± SD.

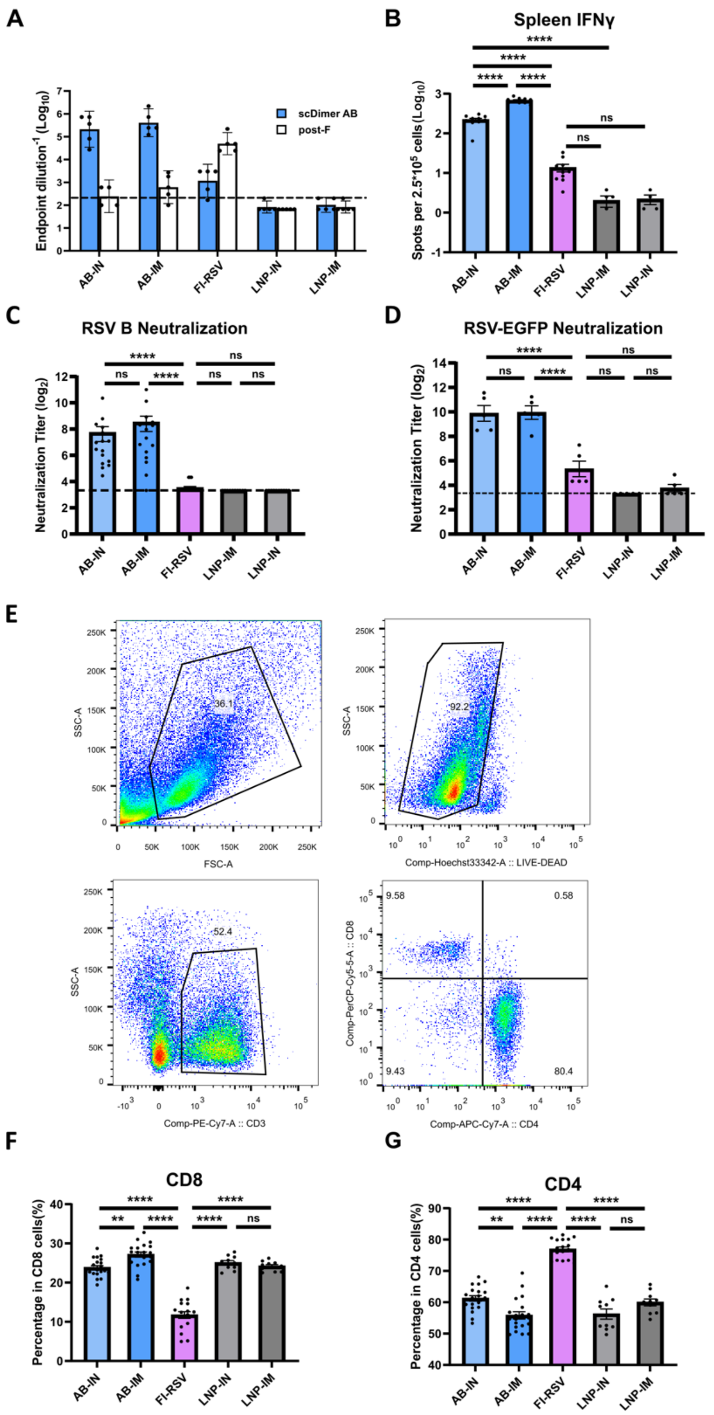


**Figure S9. Evaluations of the immune response in the immunized mice as well as those received IN or IM delivery of scDimer AB.**

(A) Endpoint titers of scDimer AB (blue) or post-F (white)-binding IgG in murine sera determined by ELISA.

(B) IFNγ-secreting spots in spleens upon re-stimulation with AB peptide pool (a mixture of monomer A and monomer B) were assessed using ELISpot on day 28.

(C) Neutralization titers in murine sera against RSV B9320 strains were determined on day 28.

(D) Neutralization titers were determined using RSV A long strain (RSV-EGFP). The serum was incubated with the virus for two days, and the fluorescence signals from RSV-EGFP were observed using CQ1 confocal image cytometer (Yokogawa).

(E) Flow cytometry gating strategy for T lymphocyte. Lung cells were obtained on 5 dpi and then stained with fluorochrome-conjugated antibodies as described in Materials and Methods. Data were collected using LSRFortessa (DB) flow cytometry machine and analyzed using FlowJo.

(F)-(G) Percentage of $CD8^+$ (F) and $CD4^+$ (G) T lymphocytes in lung on 5 dpi.

Dotted lines indicate LLD. Any value less than the LLD was assigned a value of 1⁄2 the LLD. Bars represent mean ± SD. *P* values were analyzed with One-way ANOVA (ns indicated $P > 0{\cdot}05$; * indicated $P < 0{\cdot}05$; ** indicated $P < 0{\cdot}01$, *** indicated $P < 0{\cdot}001$, **** indicated $P < 0{\cdot}0001$).
